# Plakophilin-2 Coordinates Energy Metabolism and Contractility in Cardiomyocytes, Revealing Its Roles beyond Desmosomes

**DOI:** 10.1101/2025.01.17.633239

**Authors:** J. Alex Aycinena, Anley E. Tefera, Isaac Perea-Gil, Duncan Holbrook-Smith, Kevin Williams, Reva Shenwai, Farshad Farshidfar, Brendan Ryback, Katelyn Foppe, Iris Wu, Aliya Zeng, Melissa Van Pell, Emma Xu, Joseph Woods, Samantha Jones, Yolanda Hatter, Cristina Dee-Hoskins, Jessie Madariaga, Kevin Robinson, Amara Greer-Short, Xiaomei Song, Kathryn N. Ivey, James Priest, Gretchen Argast, Timothy Hoey, Edward Driggers, Zhihong Jane Yang

## Abstract

Arrhythmogenic right ventricular cardiomyopathy (ARVC) is a familial cardiac disease associated with ventricular arrhythmias and an increased risk of sudden cardiac death. Mutations in the desmosome gene *Plakophilin-2*, *PKP2*, lead to reduction in PKP2 protein and collapse of desmosomes that is known to compromise contractility and electrical stability of cardiomyocytes. Our previous studies demonstrated the efficacy of adeno-associated virus 9 (AAV9)-mediated restoration of PKP2 expression in a cardiac specific knock-out mouse model of *Pkp2* and revealed profound changes in mRNA signatures of metabolic enzymes that were reversed by the gene replacement approach. In this study, we used PKP2-deficient mouse hearts and human induced pluripotent stem cell-derived cardiomyocytes (iPSC-CMs) to identify changes in steady-state metabolite levels associated with impaired lipid homeostasis, glycolysis, and glucose oxidation. These metabolic phenotypes align with human ARVC metabolic data and reflect an intrinsic impairment of cellular energy metabolism. Here we showed for the first time that these intracellular metabolic defects were associated specifically with poor contractility of cardiomyocytes. AAV9:PKP2 restored contractility, improved electrophysiological properties and Ca_2+_ transients. In contrast, we observed that treating PKP2-deficient cardiomyocytes pharmacologically with small molecule metabolic enhancers improved contractility but not electrophysiological properties and Ca_2+_ transients, suggesting differential sensitivity of structure-mediated functions in response to metabolic perturbance. Our study modeled and revealed a direct intracellular connection between compromised PKP2 function and metabolic impairment. We proposed that an increased risk of decoupling energy-responsive contractility from less energy-responsive electrical activities can be a new arrhythmogenic mechanism, potentially responsible for exercise-triggered cardiac adversity in ARVC disease development and progression.

## Introduction

Arrhythmogenic right ventricular cardiomyopathy (ARVC) is an inherited heart disease characterized by ventricular arrhythmias and progressive cardiac dysfunction^1,2,3,4,5^. Mutations in the desmosome gene *Plakophilin-2*, *PKP2*, result in reduction of PKP2 protein and disruption of desmosomes and gap junctions at the intercalated discs ^6,7,8^. These structural corruptions trigger a cell death response, inflammatory infiltration, and metabolic perturbation that are clinically manifested as electrical instability, cardiac structural deterioration, fibrofatty infiltration, and heart failure^9–19^. We^20^ and others^21,22,23^ reported the feasibility and the efficacy of adeno-associated virus (AAV)-mediated restoration of PKP2 in ARVC mouse models and human induced pluripotent stem cell-derived cardiomyocytes (iPSC-CMs). We found restoration of PKP2 expression through AAV9:PKP2 treatment led to a highly coordinated, durable, and dose-dependent correction of mRNA signatures of metabolic enzymes^20^. TN-401, Tenaya Therapeutics’ AAV9-based gene therapy designed to deliver fully functional human PKP2 gene, is currently at phase 1 clinical testing to treat PKP2-mutated ARVC patients.

The healthy human heart consumes fatty acids as the primary energy fuel, with the remaining sources being glucose, lactate, ketone bodies and amino acids^24,25,26^. In the failing heart, long-chain acylcarnitines, which are mitochondrial fatty acid oxidation (FAO) intermediates, are decreased^27^, accompanied with a metabolic switch from FAO to glucose-based glycolysis, as well as activation of pathways for consumption of alternative sources of energy^26,28^. Supported by clinical evidence, deranged cardiac lipid metabolism leads to intramyocardial increases of triacylglycerides (TAG) and toxic lipids that have been linked to structural myocardial damage and reduced contractility in heart failure^26,29^. These alterations in cardiac energy metabolism result in less efficient energy production in the heart and ultimately contribute to most forms of myocardial dysfunction^18,26,28,30^.

Recent advances in metabolomics and lipidomics have expanded our understanding of the mechanistic connection between energy metabolism and cardiomyopathies in human heart. Targeted metabolomics analysis revealed significant reductions in long-chain acylcarnitine and FAO, as well as increases in alternative fuel burning in heart tissues from patients with either hypertrophic cardiomyopathy (HCM) or dilated cardiomyopathy (DCM)^30,31,32^. However, myocardial TAG content was reported to show no change or decrease in the failing or nonfailing heart tissue of either HCM or DCM^31,32,33,34^.

Unlike DCM and HCM, end-stage ARVC myocardial tissue has been reported to show decreased glycolysis and pyruvate oxidation in addition to decreased long-chain acylcarnitine and FAO^18^. In addition, biopsies from the right ventricle (RV) of patients, carrying pathogenic *PKP2* variants and having >45% left ventricular ejection fraction, presented reduced glycolysis and FAO with either mRNA or protein signatures^16^. A distinct metabolic perturbation reported in ARVC is elevated intracellular lipid content, detected in heart tissue of human patients or a mouse model, and in patient-specific iPSC-CMs^35–40^. ARVC patients are subject to restriction of activity and exercise, both having been found to be associated with accelerating the disease progression and increasing the risk of life-threatening ventricular arrhythmias and sudden cardiac death^41,42,43^. Therefore, understanding how ARVC hearts handle energy challenges and metabolic stress is one of the fundamentals in assessing and managing the arrhythmogenic triggers. Specifically, we are interested in understanding the detrimental impact on cardiomyocyte energy metabolism due to PKP2 mutations and delineating the intrinsic cellular connection between PKP2-mediated structural functions and energy metabolism in the context of ARVC and poor cardiomyocyte contractility and electrical instability.

In broadly interrogating ARVC-associated metabolic changes, we compared nontargeted metabolomics profiles of mouse heart tissue from cardiac-specific knock-out, *Pkp2-cKO*, with healthy mouse heart tissue and also with AAV9:PKP2 treated *Pkp2-cKO* mouse heart tissue, to identify altered metabolic pathways that responded to AAV9:PKP2-mediated restoration of PKP2 expression. Functionally, we investigated how PKP2 mutation-associated alterations in bioenergetics affected contractility, electrophysiological properties, and Ca_2+_ transients using human iPSC-CMs. Both model systems confirmed altered lipid homeostasis, glycolysis/glucose oxidation that agreed with the human data^18^. We reported here that PKP2 deficiency led to impaired energy production correlating with poor contractility of cardiomyocytes. Treatment with AAV9:PKP2 restored contractility, electrophysiological properties, and Ca_2+_ transients, whereas treating PKP2-deficient cardiomyocytes with metabolic enhancers improved contractile function but not electrophysiological properties or Ca_2+_ transients. These results suggested structure-mediated functions, such as contractility, electrophysiological properties, and Ca_2+_ transients, differentially responded to metabolic perturbance, and decoupling energy-responsive contractility from the less responsive electrophysiological properties and Ca_2+_ transients may be an arrhythmogenic substrate occurring in the presence of energy challenge and metabolic stress. We concluded that PKP2 coordinates energy metabolism and contractility in the cardiomyocytes and that compromised PKP2 led to metabolic impairment and cardiomyocyte dysfunctions at the intracellular level.

## Methods

More detailed materials and methods are available in the Supplementary Material.

### Animal studies

Animal studies were performed according to Tenaya Therapeutics’ animal use guidelines. The animal protocols were approved by the Institutional Animal Care and Use Committee (IACUC number: 2020.007).

### Mouse model and route of AAV administration

Tenaya licensed a cardiomyocyte-specific, tamoxifen-activated *Pkp2-cKO (αMyHC-Cre-ER(T2)/Pkp2_fl/fl_)* mouse line in the C57BL/6 background from Dr. Mario Delmar, NYU Grossman School of Medicine^13^. The *Pkp2-cKO* animals were induced with tamoxifen at 0.1mg/g for 3 consecutive days. Tamoxifen injection activates Cre recombinase in Cre-positive animals and induces homozygous deletion of *Pkp2* gene. TN-401 or AAV9:mouse Pkp2 (AAV9:mPkp2) was given as a single dose via retro-orbital injection before or after animals were induced with tamoxifen.

### Untargeted polar metabolome profiling

Polar metabolites were extracted from mouse heart tissue and human iPSC-CM culture. Network-based integration of metabolomics and transcriptomics data were performed and detailed in the Supplementary Material.

## Results

### AAV9:PKP2 largely maintained key mRNA signatures of cardiac energy metabolism that were perturbed in *Pkp2-cKO* ARVC mouse heart

Our published work showed that a single dose of AAV9:mPkp2 treatment in *Pkp2-cKO* ARVC mice after overt cardiomyopathy halted disease progression and extended median lifespan by ≥ 50 weeks post induction of *Pkp2* deletion^20^. RNA sequencing was carried out for heart tissues collected from either the therapeutic mode of treatment (virus injection after overt structural change) or the preventive mode of treatment (virus injection before overt structural change) (Fig. 1a). When comparing vehicle-treated *Pkp2-cKO* animals vs WT, the top two negatively enriched gene sets identified by Gene Set Enrichment Analysis (GSEA)^44^ were mitochondrial dysfunction and cardiac muscle contraction, suggesting that PKP2 deficiency led to impaired oxidative phosphorylation and contractile functions. Transcriptional changes in key enzymes involved in both oxidative and glycolytic metabolic pathways were observed in heart failure and cardiomyopathies^18,26,28,30,32^. To investigate a connection between PKP2 and cardiac energy metabolism, we categorized mRNA changes in major cardiac metabolic pathways of lipid homeostasis and glycolysis/glucose oxidation with highlighted enzymes as known points of regulation (Fig. 1b). Glucose metabolism generates ATP from cytoplasmic glycolysis and the mitochondrial oxidation of the pyruvate derived from glycolysis. Fatty acid homeostasis is further categorized to gene classes of FAO, TAG dynamics, and fatty acid synthesis (Fig. 1c). In addition, mRNA changes in master regulators of cardiac metabolism, members of the peroxisome proliferator-activated receptor (PPAR) family^45^, were included (Fig. 1c).

**Figure 1.**
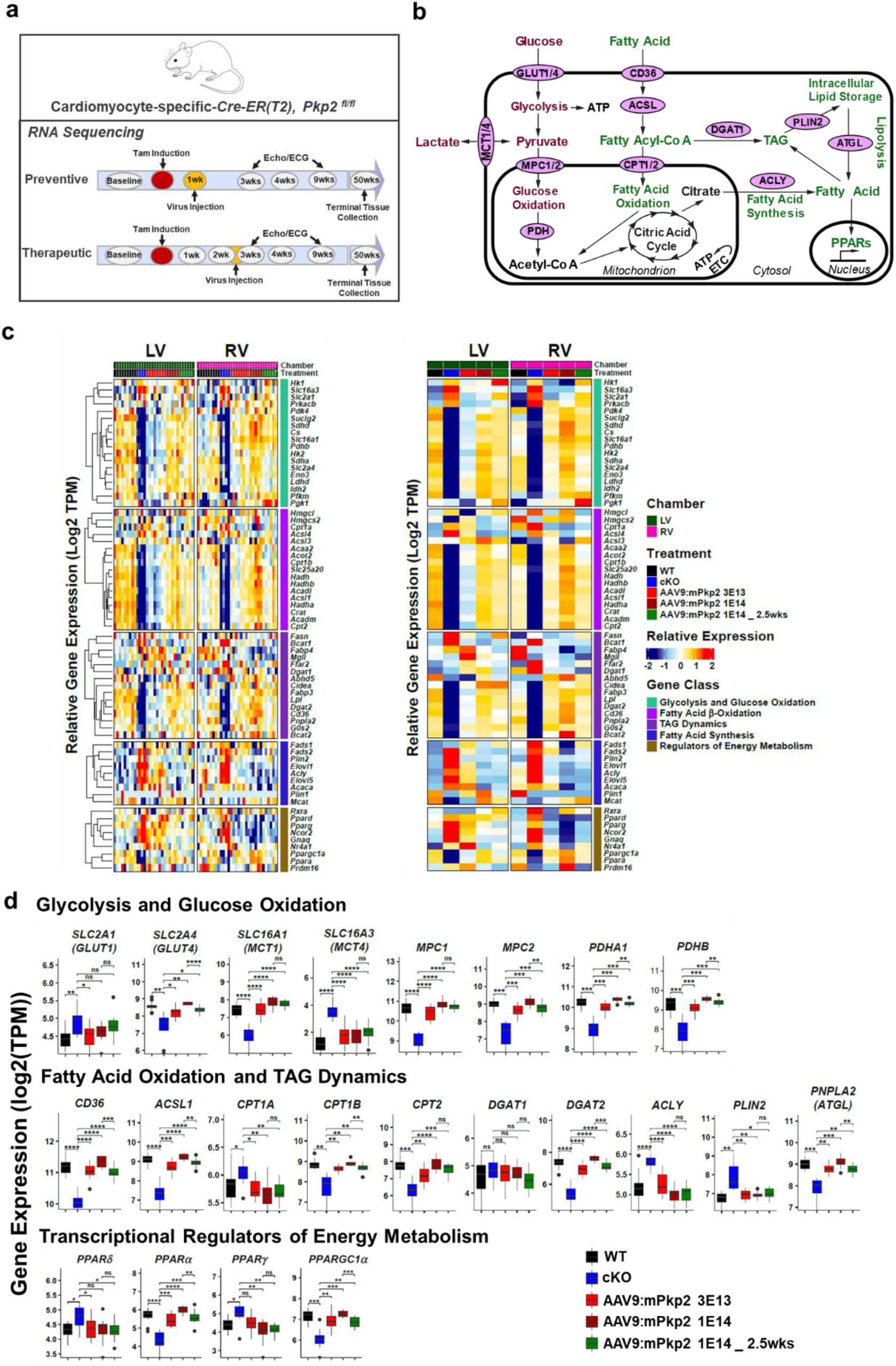
AAV9:PKP2 largely maintained key mRNAsignatures of cardiac energy metabolism that was perturbed in *Pkp2-cKO* ARVC mouse model. **a,** Study design to evaluate AAV9:PKP2 efficacy at 51 weeks post tamoxifen induction of *Pkp2* deletion in *Pkp2-cKO* ARVC mouse model. AAV9:mPkp2 was dosed at 3E13 and 1E14 vg/kg either 1 week after the induction (the preventive mode of treatment) or at 1E14 vg/kg at 2.5 weeks after induction (the therapeutic mode of treatment). **b,** Two key cardiac energy productions, glucose oxidation and fatty acid b-oxidation was highlighted with enzymes of known points of regulation. Additional mRNA analyses were carried out to examine triacylglycerol (TAG) dynamics, fatty acid synthesis, and expression of PPAR transcriptional regulators of energy metabolism. **c,** Relative gene expression of selected genes was measured by RNA-seq. Samples were sorted by RV (right ventricle) and LV (left ventricle) chambers and treatment groups. Genes were categorized by gene classes. Left panel shows heatmap of gene expression of each individual animal and its corresponding chambers presented in scaled log2-transformed. Right panel depicts a scaled average across samples of each treatment group. Number of animals in each treatment group used for RNA sequencing were 9 WT, 4 cKO, 8 cKO + AAV9:mPkp2 at 3E13 vg/kg, 5 cKO + AAV9:mPkp2 at 1E14 vg/kg (the preventive mode), and 6 cKO + AAV9:mPkp2 at 1E14 vg/kg (the therapeutic mode) with both RV and LV collected. **d,** Boxplots showed group-wise gene expression of the selected gene classes. Each box showed the distribution of expression values in the following manner: the midline represented the median expression value, the box indicated the interquartile range where the middle 50% of values lie, and the whiskers at the top and bottom of each box represented the range of values outside the interquartile range. The black dots represent values that fall outside the 2_nd_ and 3_rd_ quartiles. Values were log 2 of TPM (Transcripts Per Million) and were aggregated from LV and RV. Comparison p values were calculated by Student’s t-test: p values: *p<0.05, **p<0.01, ***p<0.001****p<0.0001.

In agreement with the published human ARVC tissue data^18^, we observed mRNA signatures predominantly showing depressed FAO and glycolysis/glucose oxidation and increased fatty acid synthesis in both right and left ventricles of *Pkp2-cKO* mouse heart (Fig. 1c and 1d). In addition, mRNA signatures of TAG dynamics were greatly perturbed with respect to storage and mobilization via lipolysis. PPARγ activation has been associated with promoting fatty acid uptake, TAG formation, and storage in lipid droplets^45^. PPARγ mRNA was significantly upregulated in contrast to decreased PPARα mRNA. In both preventive and therapeutic modes of AAV9:mPkp2 treatment, we observed significant long-term reversal of the disturbed metabolic mRNA signatures with the preventive mode supporting the most complete reversal among age-matched animals (Fig. 1c and 1d). Similar conclusions were drawn based on a short-term study where metabolic reversal revealed a dose-dependent response to TN-401 expression level, suggesting a direct connection between PKP2 function and the cardiac energy metabolism (Supplemental Fig. 1).

### Integrated metabolomics and transcriptomics identified metabolic impairment in *Pkp2-cKO* cardiac tissue that was reversed by AAV9:PKP2

We used the *Pkp2-cKO* ARVC mouse model to assess the feasibility and the efficacy of TN-401 on reversing the metabolic impairment^20^. Nontargeted polar metabolomics using FIA-TOF mass spectrometry^46^ were conducted on cardiac tissues collected from either the therapeutic or the preventive mode of treatment (the therapeutic mode in Fig. 2a and the preventive mode in Supplementary Fig. 2a). Integrated nontargeted polar metabolomics and RNA sequencing data^47,48^ were used to enhance confidence of detection as significantly changed steady-state metabolites in nodes and genes in rods were clustered and color-coded based on metabolic pathways in response to AAV9:PKP2, AAV9:mPkp2 or TN-401, treatment (Fig. 2b and Supplementary Fig. 2b). Metabolites and associated enzymes were highlighted for FAO and glycolysis and glucose oxidation. In addition, their corresponding sequencing reads of connecting genes highlighted on the cluster were summarized in box plots (Fig. 2b). The significance and fold change of a set of metabolites identified by the integrated omics were represented in a heatmap (Fig. 2c). The integrated metabo-transcriptional network and multi-omics analysis of individual transcripts identified reduced abundance of acylcarnitines in the pathway ‘Fatty Acid β-Oxidation’ in cKO mice which was reversed by TN-401 treatment as is shown in the heatmap. The impaired FAO was further supported by mRNA reduction in CPT1B (carnitine palmitoyltransferase 1B), CACT (SLC25A20, carnitine-acylcarnitine translocase), CRAT (carnitine O-acetyltransferase), and ACSL (acyl-CoA synthetase long chain family member) and their reversal by AAV9:mPkp2 (box plots, Fig. 2b). Our data agreed with prior observations of protein reduction for CPT1B, CACT, CRAT, and ACSL, and the corresponding reduction of long-chain acylcarnitines in human hearts with end-stage ARVC^18^. Not reported by prior human data, we observed that ATGL mRNA (PNPLA2, adipose triglyceride lipase) was significantly down-regulated in cKO heart and reversed by AAV9:PKP2, suggesting a reduction in intracellular TAG-derived fatty acid mobilization in the context of unameliorated disease (Fig. 1b, 1d, and 2b).

**Figure 2.**
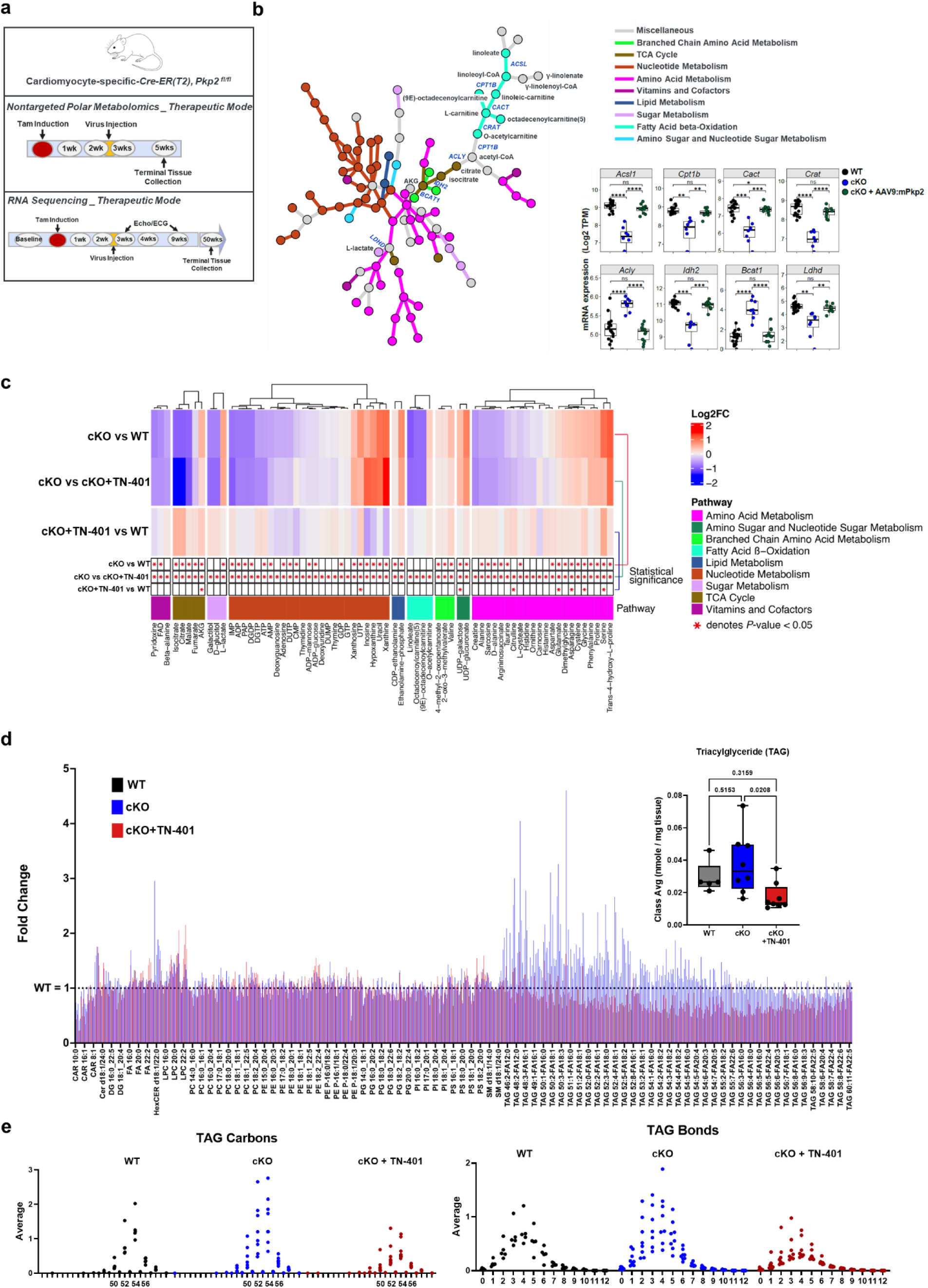
Integrated metabolomics and transcriptomics identified metabolic impairment in *Pkp2-cKO* cardiac tissue that was reversed by AAV9:PKP2. **a,** Multiomic study design for nontargeted polar metabolomics and RNA sequencing analysis, respectively, to evaluate AAV9:PKP2 efficacy in *Pkp2-cKO* ARVC mouse model. TN-401 was dosed at 1E14 vg/kg at 2.5 weeks after tamoxifen induction of *Pkp2* gene deletion (the therapeutic mode of treatment) and heart tissue was collected 5 weeks after tamoxifen induction and 2.5 weeks after virus injection. FIA-TOF mass spectrometry was used for nontargeted metabolomics. RNA sequencing analysis used the dataset of the therapeutic mode as shown in Figure 1. **b,** Cluster presentation of integrative metabolomics and RNA sequencing showed significantly changed metabolites in nodes and genes in rods in response to AAV9:PKP2 in the therapeutic mode of treatment when compared to the untreated *Pkp2-cKO* heart tissue. Metabolites and genes were categorized by functional pathways with highlights in fatty acid homeostasis, TCA cycle, and glycolysis. Boxplots showed expression of significantly changed genes highlighted on the cluster and measured by RNA-seq. Each box showed the distribution of expression values in the following manner: the midline represented the median expression value, the box indicated the interquartile range where the middle 50% of values lie, and the whiskers at the top and bottom of each box represented the range of values outside the interquartile range. The black dots represent values that fall outside the 2_nd_ and 3_rd_ quartiles. Values were log 2 of TPM (Transcripts Per Million) and were aggregated from LV and RV. Comparison p values were calculated by Student’s t-test: p values: *p<0.05, **p<0.01, ***p<0.001, ****p<0.0001. **c,** Heatmap showed mean values of log2 fold change of metabolites with each column representing cKO vs WT, cKO vs TN-401 treated, and TN-401 treated vs WT, respectively. Metabolites were categorized by functional pathways. **d,** Shotgun lipidomics quantitatively targeted >1400 lipid species across 19 lipid subclasses using standards and Sciex QTRAP 5500 system with a differential mobility spectrometry (DMS) (SelexION), yielding 981 reported lipid species normalized to wet tissue mass (nmoles/mg). Lipidomics data was subsequently filtered by data occupancy so only lipid species reported in ≥75% of tissue samples per treatment group were analyzed. After filtering for data occupancy, 691 lipid species from 17 subclasses were analyzed. Lipid species were shown as the average fold change of cKO or TN-401 treated vs WT. Inset box plot is the triacylglyceride class average in cardiac tissue of WT, cKO, and TN-401 treated. Black dots represent the class average from each animal, while the midline represents the median of all animals, and the box indicates the interquartile range where the middle 50% of values lie, and the whiskers at the top and bottom of each box represent the range of values outside the interquartile range. Statistical evaluation was performed using ordinary One-Way ANOVA (Tukey’s post-hoc test). **e,** All bulk triacylglycerol species containing any given numbers of carbon or double bonds were averaged and profiled in cardiac tissue of WT, cKO, and TN-401 treated. Number of animals in each treatment group used for RNA sequencing were 9 WT, 4 cKO, and 6 cKO treated by AAV9:mPkp2 at 1E14 vg/kg (the therapeutic mode). Number of animals in each treatment group used for metabolomics were 9 WT, 8 cKO, 8 cKO treated by TN-401 at 1E14 vg/kg (the therapeutic mode), and 5 WT, 8 cKO, 8 cKO treated by TN-401 at 1E14 vg/kg (the preventive mode).

Furthermore, in agreement with reduced coupling of glycolysis with pyruvate oxidation in human ARVC heart, the cKO heart showed increased L-lactate levels (Fig. 2c) and reduced mRNA of PFKM (phosphofructokinase, muscle) and LDHD (lactate dehydrogenase D) (Fig. 1c), suggesting less pyruvate generated and/or less lactate was oxidized to pyruvate. Impaired TCA (tricarboxylic acid) cycle was demonstrated by reduced metabolomics signal intensity for citrate, malate, and fumarate, as well as increased AKG (α-ketoglutarate) with correspondingly reduced mRNA of IDH2 (isocitrate dehydrogenase 2) and BCAT1 (branched chain amino acid transaminase 1) (Fig. 2b).

Perturbed energy metabolism of the ARVC mouse model, particularly impaired FAO and increased *de novo* lipid synthesis, was further investigated by shotgun lipidomics^49,50^ to confirm increased TAG levels, which were normalized in TN-401 treated cKO heart (Fig. 2d). Detailed analyses of all TAG species showed specific increases in TAGs with 50-56 carbon numbers and 2-5 double-bonds (Fig. 2e). Lipidomics profiling of 685 patient plasma samples identified increased lipid species with a low carbon number and double-bond content, for example, TAG (54:2), to predict risk of cardiovascular disease in the prospective population-based Bruneck study^51^.

Additional integrated omics were performed between metabolomics and RNA sequencing in response to a shorter-term treatment of TN-401 in the preventive mode (Supplementary Fig. 2). Intriguingly, the significantly changed metabolites identified in the preventive mode of treatment showed a partial overlap with the metabolites identified in therapeutic mode. This observation suggested that reversal of metabolic impairment before and after overt structural changes by AAV9:PKP2, AAV9:mPkp2 or TN-401, likely depends on the extent of the impairment and the metabolic profiling at a given intervention time reflects a snapshot of metabolic changes.

As integrated omics were conducted based on the metabolic profiling of the heterogenous cardiac tissue, we moved the functional studies to iPSC-CMs to specifically identify intrinsic connections of PKP2 to energy metabolism at the cellular level.

### Acute PKP2 silencing disrupted lipid homeostasis and led to increased intracellular lipid granules in iPSC-CMs

A highly regulated balance between fatty acid uptake, TAG dynamics, and FAO ensures effective fatty acid energy metabolism and proper heart function^26^. Resembling adult-like cardiac metabolism^52^, we examined intracellular fatty acid homeostasis, specifically fatty acid uptake, fatty acid storage, and fatty acid mobilization from the TAG pool in response to acute PKP2 silencing and other metabolic modulators in iPSC-CMs (Fig. 3a).

**Figure 3.**
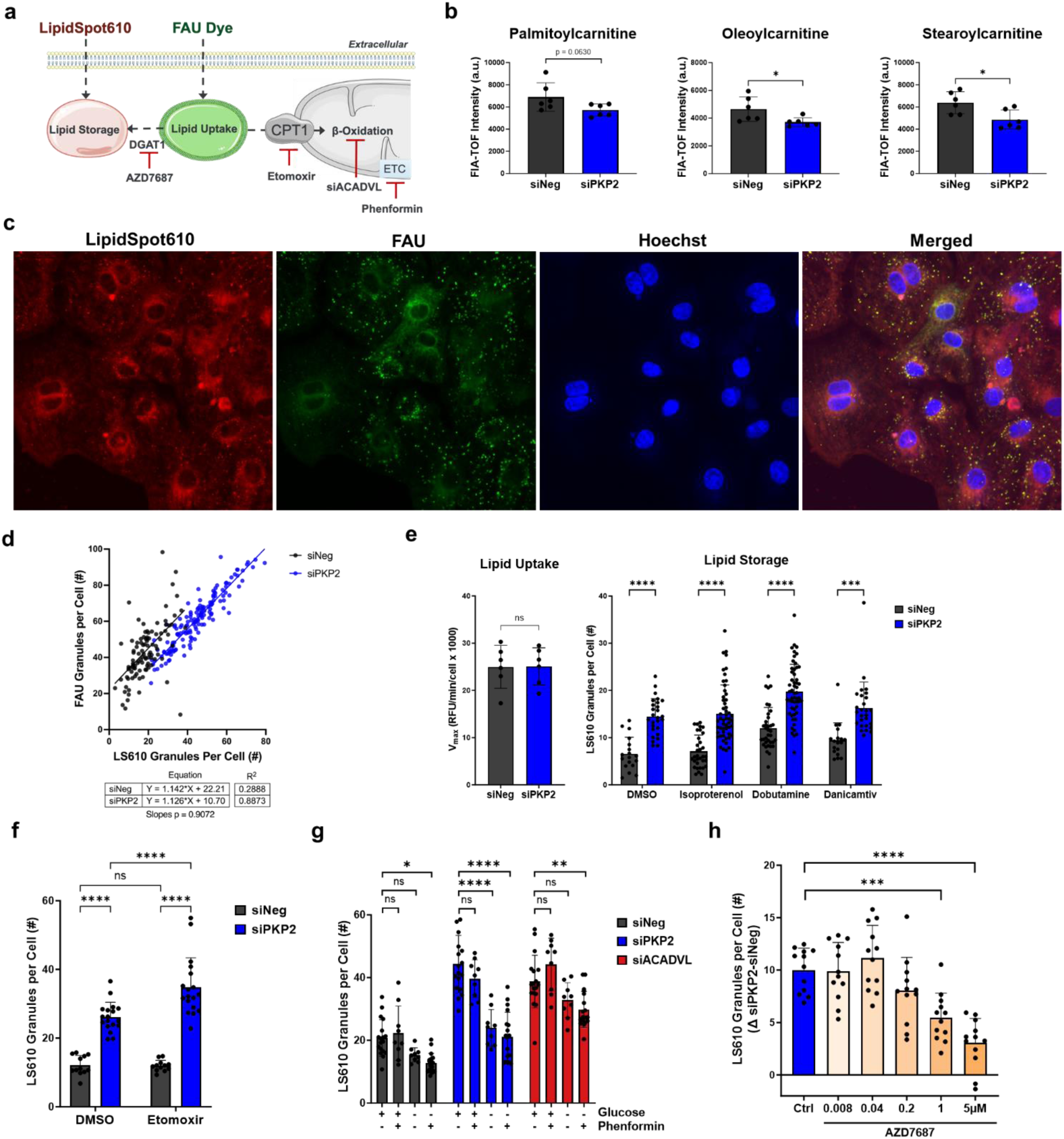
Acute PKP2 silencing disrupted lipid homeostasis and led to increased intracellular lipid granules in iPSC cardiomyocytes. **a,** Study design to evaluate lipid homeostasis in response to acute PKP2 silencing and metabolic modulators. **b,** FIA-TOF mass spectrometry was used to measure acylcarnitines in response to acute depletion of PKP2 (n = 6 technical replicates, 3 wells of 12-well plate were pooled as 1 technical replicate). The negative control for siRNA-mediated silencing is siNeg. **c,** Confocal imaging using two lipid dyes, QBT Fatty Acid Uptake (Molecular Devices) for measuring real-time fatty acid uptake (FAU) and LipidSpot 610 for measuring fatty acid storage in fixed iPSC cardiomyocytes after performing FAU. FAU dye in green, storage dye in red, and nuclei in blue. **d,** Immunofluorescence post cell fixation showed co-stained of FAU dye and LipidSpot610 with more cells stained by LipidSpot610 in siPKP2 than siNeg. **e,** Maximum FAU rate, V_max_ (Relative Fluorescence Units/minute/cell x 1000), did not change in response to acute silencing of PKP2. Fatty acid storage was increased in response to acute silencing of PKP2 and was not alleviated by 48-hour treatment of isoproterenol, dobutamine, or danicamtiv, respectively. QBT Fatty Acid Uptake: n = 6 technical replicates; LipidSpot 610: n = 18-54 technical replicates. **f,** Inhibition of β-oxidation by etomoxir increased intracellular lipid granules in response to PKP2 acute silencing (n = 9-18 technical replicates). **g,** Comparing to acute silencing of ACADVL, siPKP2 showed a similar extent of increased lipid granules (n = 9-18 technical replicates). Glucose starvation reduced lipid granules. Inhibition of mitochondrial complex 1 by 40µM phenformin did not increase lipid granules with or without glucose starvation. **h,** Inhibition of DGAT1 by AZD7687 reduced amount of lipid granules in response to PKP2 deficiency, suggesting reduction of TAG formation (n = 12 technical replicates). Data represented as mean ± SD. Comparison p values were calculated by Prism unpaired *t*-test for acylcarnitine levels and V_max_, ordinary One-Way ANOVA (Tukey’s post-hoc test) for all other comparisons: p values: *p<0.05, **p<0.01, ***p<0.001, ****p<0.0001.

First, nontargeted polar metabolomics showed a trend towards reduced long-chain acylcarnitines, palmitoylcarnitine, oleoylcarnitine, and stearoylcarnitine, in iPSC-CMs in response to acute silencing of PKP2 (Fig. 3b). Correspondingly, a real-time fluorescence-based lipid uptake assay was used to analyze fatty acid uptake (FAU, green) kinetics and a fixed-cell based lipid droplet dye (LS610, red) to quantify lipid storage. After performing real-time FAU assay, iPSC-CMs were fixed and both green and red channels were imaged (Fig. 3c) and quantified by granule counts (Fig. 3d).

Acute silencing of PKP2 led to an upward shift in the total number of granules positively stained with LS610 within the quantified microscopic fields of view (Fig. 3d). Real-time kinetic readouts on the maximum FAU rate normalized to cell counts did not respond to acute PKP2 silencing (Fig. 3e, left panel). In addition, short-term treatment of iPSC-CMs with small molecules such as isoproterenol (non-selective β-adrenergic agonist), dobutamine (β1-adrenergic agonist), and danicamtiv (cardiac myosin activator) did not alleviate increased lipid granules in response to acute silencing of PKP2 (Fig. 3e, right panel).

The nature of these increased lipid granules in the presence of PKP2 depletion was further investigated by small-molecule metabolic modulators. Inhibition of the oxidation of long-chain fatty acids within mitochondria by etomoxir, an irreversible inhibitor of CPT1, further increased the abundance of intracellular lipid granules (Fig. 3f). The extent of increased lipid granules in response to acute silencing of PKP2 was directly compared to acute silencing of ACADVL, encoding very long-chain specific acyl-CoA dehydrogenase that catalyzes the first step of FAO in mitochondria (Fig. 3g). Increased lipid granules observed by either depleting PKP2 or ACADVL can be alleviated by glucose starvation, putatively via increasing fatty acid mobilization from TAG storage to fuel FAO (Fig. 3g). Interestingly, phenformin treatment inhibited mitochondrial complex 1^53^ and did not exacerbate lipid granules (Fig. 3g). We observed that treatment of granule-rich PKP2-depleted cells with a highly selective DGAT1 (diacylglycerol acyltransferase 1) inhibitor, AZD7687^54^, which is known to impede TAG formation, reduced granule abundance in a dose-dependent manner (Fig. 3h), providing strong-mechanism-based evidence that TAGs comprised a significant portion of the granules. Therefore, the presence of increased TAGs observed in *Pkp2-cKO* mouse heart tissue was recapitulated in the form of increased intracellular TAG-rich granules in human iPSC-CMs in response to acute PKP2 depletion.

### Acute PKP2 silencing decreased oxidative and glycolytic metabolism resulting in decreased contractility in human iPSC-CMs

Both mouse heart tissue and human iPSC-CMs revealed profound impairment in energy metabolism, particularly disrupted lipid homeostasis, in response to PKP2 deficiency. We therefore examined how impaired energy metabolism altered tissue or cellular fuel choices, with the potential to impact contractile function of human iPSC-CMs (Fig. 4a).

**Figure 4.**
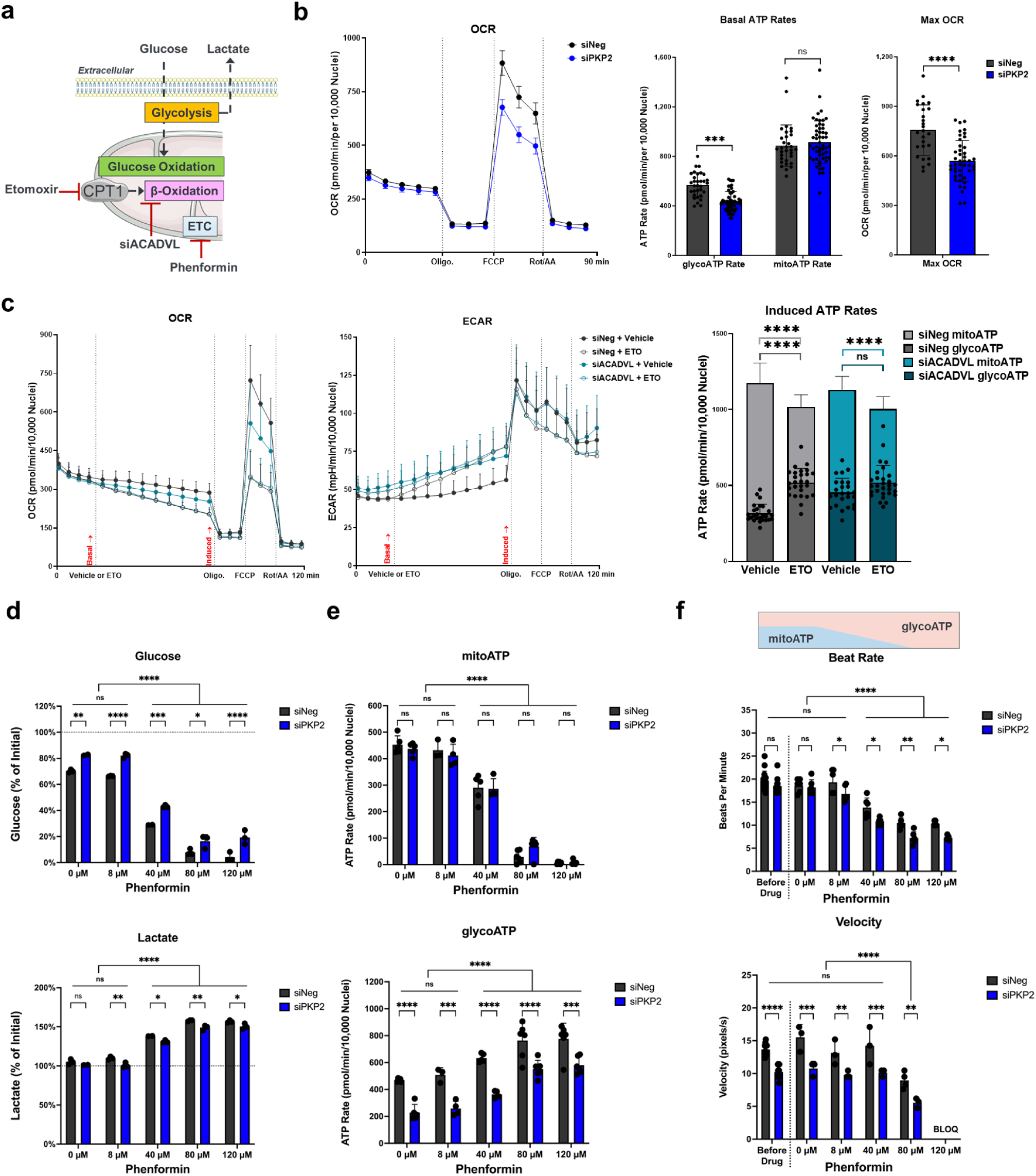
Acute PKP2 silencing decreased oxidative and glycolytic metabolism resulting in decreased contractility in iPSC-CMs. **a,** Study design to evaluate the metabolic status of human iPSC cardiomyocytes in response to acute silencing of PKP2 and metabolic modulators. **b,** Acute silencing of PKP2 significantly decreased glycolytic and oxidative metabolism as indicated by decreased glycoATP rate and Max OCR determined by Seahorse MitoStress test (n = 25-53 technical replicates). Comparison p values were performed by Prism: basal ATP rates by ordinary One-Way ANOVA (Tukey’s post-hoc test), ***p<0.001, and Max OCR by unpaired t-test, ****p<0.0001. **c,** Inhibition of β-oxidation by etomoxir or silencing ACADVL decreased fatty acid oxidative metabolism and increased glycolysis determined by Seahorse MitoStress test (n = 24-27 technical replicates). Left kinetic curves indicated where basal or vehicle/etomoxir-induced ATP rates were calculated. Comparison p values were calculated by ordinary One-Way ANOVA (Tukey’s post-hoc test). **d,** Percent of the initial amount of glucose or lactate in the culturing media was measured by Bioprofile Flex2 in response to acute silencing of PKP2 and a range of phenformin treatment (n = 2-3 technical replicates, 3 wells of 96-well plate were pooled as 1 technical replicate at both initial time point and final assay time point). **e,** MitoATP and glycoATP rates were determined in response to acute silencing of PKP2 and a range of overnight phenformin treatment in the presence of 1 mM pyruvate (n = 3-6 technical replicates per treatment). Inhibition of mitochondrial complex 1 by phenformin switched from oxidative phosphorylation to glycolysis as represented by decrease mitoATP and increased glycoATP. **f,** Contractility in response to acute silencing of PKP2 and a range of overnight phenformin treatment was evaluated using SONY live cell imaging and Pulse video analysis (n = 6-16 technical replicates per treatment) (Curi Bio)_55_. Switching from oxidative metabolism to glycolysis supported poor contractility as measured by beat rate and contraction velocity. Average nuclear counts from live cells were used to normalize contraction velocity. Phenformin treatment at 120 µM inhibited cell contraction such that velocity was below limit of quantification (BLOQ). Comparison p values shown on **d**, **e**, and **f** were calculated by ordinary Two-Way ANOVA (Sidak’s post-hoc test): p values: *p<0.05, **p<0.01, ***p<0.001, ****p<0.0001.

We used Seahorse-based bioenergetic analysis to evaluate the glycolytic and oxidative phosphorylation state of iPSC-CM in response to acute PKP2 silencing. PKP2 silencing led to a significant reduction in the glycolysis-derived ATP production rate (“glycoATP”). Although the basal mitochondrial respiration-derived ATP production rate (“mitoATP”) was not affected, maximal oxygen consumption rate (Max OCR) was significantly decreased, confirming impaired glycolytic and oxidative metabolism due to PKP2 silencing (Fig. 4b). However, in the context of FAO inhibition by either etomoxir or ACADVL silencing (Fig. 4c), bioenergetic analysis showed a 13 or 11% reduction in mitoATP rate, respectively, and a 39 or 13% increase in glycoATP rate, respectively (Fig. 4c, the bar graph only showed induced ATP rate). In contrast to impaired oxidative and glycolytic metabolism due to PKP2 deficiency (Fig. 4b), FAO reduction by etomoxir and ACADVL silencing in iPSC-CMs having wild-type PKP2 (Fig. 4c) led to fuel switch to glycolysis.

Direct measurements of glucose and lactate in the culturing media further confirmed impaired glycolysis with less glucose consumption and a concomitant trending reduction of lactate produced in response to PKP2 silencing as compared to the control, siNeg (at 0 µM phenformin, Fig. 4d). Increasing phenformin, an inhibitor of mitochondrial complex 1, increased glucose usage by tuning down mitochondrial oxidative phosphorylation, and was accompanied with increased lactate and enhanced glycolysis (at 8-120 µM phenformin, Fig. 4d). PKP2 silenced cardiomyocytes showed no change in mitoATP and a lower glycoATP rate (at 0 µM phenformin, Fig. 4e) and in the presence of increasing phenformin, kept no change in mitoATP rates and lower glycoATP rates than the control (at 8-120 µM phenformin, Fig. 4e). Phenformin titration showed that iPSC-CMs switched to glycolysis as oxidative phosphorylation was suppressed, and that glycolytic metabolism alone (>80µM phenformin) supported poor contractile function as measured by reduced beat rate and contraction velocity, both being worse in PKP2 silenced cardiomyocytes (Fig. 4f).

It is worth noting that the polar metabolomics of cKO heart tissue (Fig. 2b and 2c) and PKP2-slenced iPSC-CMs (Fig. 4d) revealed opposite effects on lactate levels, with steady-state lactate levels increased in the cKO context but lowered in iPSC-CMs. This apparent discrepancy is consistent with reduction of oxidative mitochondrial metabolism of pyruvate in each system, likely reflecting the documented uptake of circulating lactate by cardiac tissue^25,28^, a fuel source that is not available to the iPSC-CMs grown in culture. The reduced oxidative metabolism of circulating lactate possibly explained increased lactate levels in the cKO heart tissue whereas reduced glycolysis in iPSC-CMs was an intrinsic cellular response to PKP2 silencing. Thus, this discrepancy was seen to support the shared mechanism of impaired glycolysis and glucose oxidation between cKO mouse heart and human iPSC-CMs. At the cell level, our results also supported that residual glycolytic metabolism was not the optimal energy source to propel contractile function of cardiomyocytes and worsened when desmosome function was corrupted due to PKP2 depletion.

### Isogenic human iPSC-CMs carrying a pathogenic PKP2 mutation showed impaired oxidative and glycolytic metabolism and contractile function

Acute depletion of PKP2 helped us understand a ‘rapid’ metabolic response that can be attributed to the metabolic flexibility observed in cardiomyocytes. To understand a longer-term or stable PKP2-associated metabolic alterations that leads to metabolic impairment, we generated heterozygous, PKP2_Het_, and homozygous, PKP2_Hom_, isogenic iPSC-CMs carrying a pathogenic mutation c.2146G>C^56^. Differentiation of wild-type, PKP2_Het_ and PKP2_Hom_ isogenic cardiomyocytes was confirmed by cardiac troponin T expression (Supplementary Fig. 3a). PKP2 and two other desmosome proteins, desmoplakin (DSP) and plakoglobin (PKG), were quantified based on signal intensity of immunofluorescence (Supplementary Fig. 3a and 3b), supporting the observation of a stoichiometric relationship between PKP2 with other desmosome components^20^.

Functional impact by PKP2 genotypes, PKP2_Het_ and PKP2_Hom_, was characterized by contractility (Supplementary Fig. 3c), electrophysiological properties (Supplementary Fig. 3d), and levels of long-chain acylcarnitines (Supplementary Fig. 3e), confirming that PKP2 dosage effects between isogenic PKP2_Het_ and PKP2_Hom_ iPSC-CMs were associated with most functional phenotypes that we characterized. We wanted to emphasize that the reversibility of metabolic impairment and the subsequent improvement of functional outcomes, for example, contractility, electrophysiological properties, and calcium transient, should be delineated in PKP2_Hom_ isogenic cells. PKP2_Het_ cardiomyocytes, expressing some PKP2, could confound data interpretation when these structural functions are canonical PKP2 functions.

Increased lipid granules were observed in PKP2_Hom_ isogenic mutant cells relative to the WT isogenic control (Fig. 5a). Nontargeted polar metabolomics confirmed the same observation seen in the acute PKP2 silencing, that long-chain acylcarnitines, palmitoylcarnitine, oleoylcarnitine, and stearoylcarnitine, were reduced in isogenic PKP2_Hom_ cells (Fig. 5b). Bioenergetics analyses of isogenic mutant cells showed significantly impaired oxidative and glycolytic metabolism as demonstrated by reductions in basal and vehicle/etomoxir-induced glycoATP and mitoATP, and Max OCR (Fig. 5c). In addition, Max OCR was affected less by etomoxir treatment in PKP2_Hom_ isogenic cells, suggesting a limited capacity to switch to glycolysis as a result of PKP2 deficiency (Fig. 5c).

**Figure 5.**
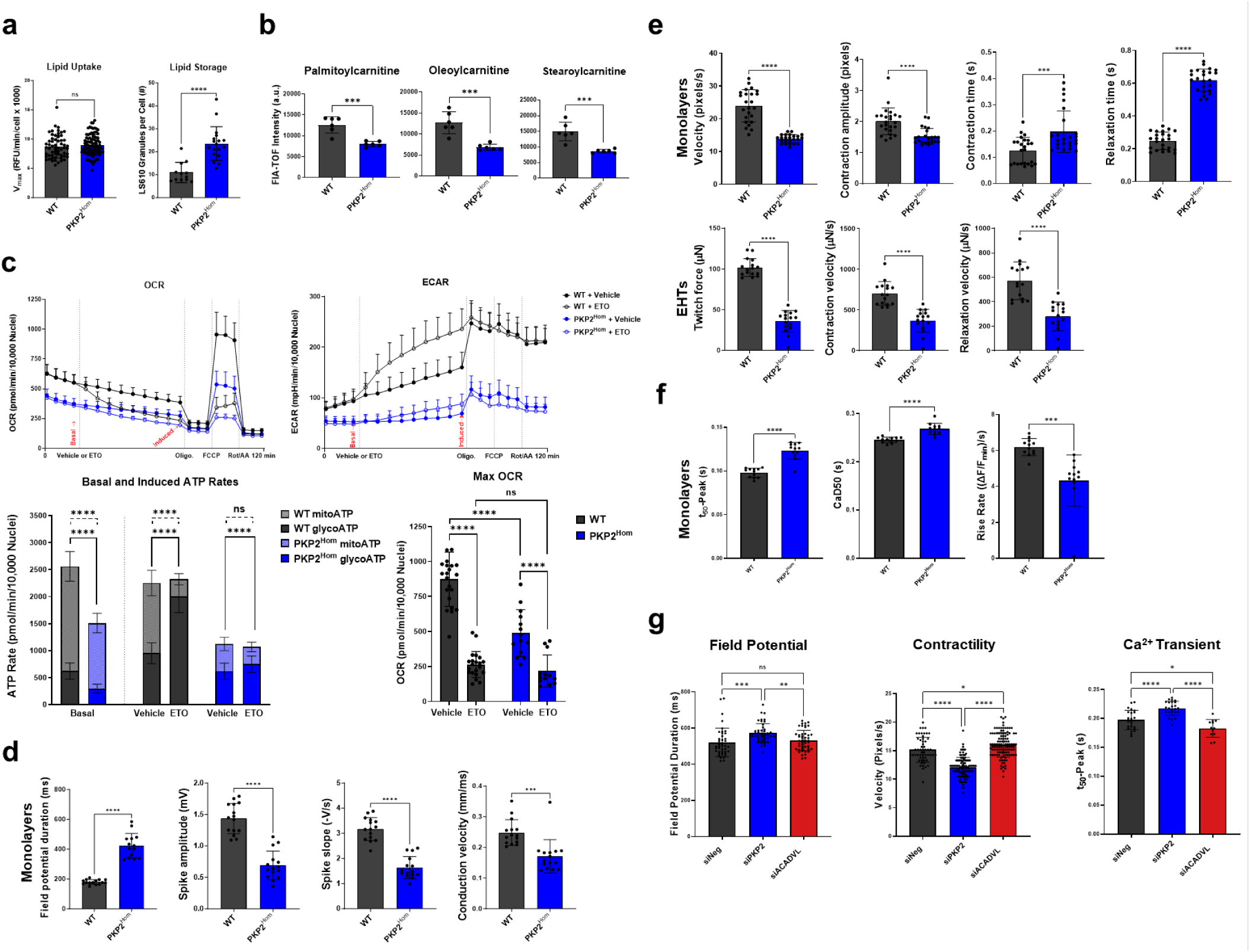
Isogenic human iPSC cardiomyocytes carrying a pathogenic PKP2 mutation showed impaired oxidative and glycolytic metabolism and contractile function. **a,** PKP2_Hom_ showed increase in lipid granules and no change in lipid uptake compared to WT isogenic cell quantified by confocal imaging using two lipid dyes, QBT Fatty Acid Uptake (Molecular Devices) for measuring real-time fatty acid uptake (FAU) (n = 59-90 technical replicates) and LipidSpot 610 for measuring fatty acid storage in fixed iPSC cardiomyocytes after performing FAU (n = 12-18 technical replicates). **b,** FIA-TOF mass spectrometry measurement showed reduced acylcarnitines in PKP2_Hom_ compared to WT isogenic cells (n = 6 technical replicates, 3 wells of 12-well plate were pooled as 1 technical replicate). **c,** PKP2_Hom_ showed reduced glycolytic and oxidative metabolism evaluated by the Seahorse MitoStress test (n = 14-43 technical replicates). Top kinetic curves indicated where basal or vehicle/etomoxir-induced ATP rates were calculated. **d,** Monolayers MEA parameters showed prolonged field potential duration (FPD) along with depressed spike amplitude, conduction velocity, and spike slope using the Maestro Pro Microelectrode array (MEA) platform (Axion Biosystems)_58_ (n = 15 technical replicates). **e,** Monolayers (n = 24 technical replicates) and engineered heart tissues (EHTs) (n = 16 technical replicates) showed reduced contraction force and velocity, and prolonged contraction and relaxation time in the mutant lines using SONY live cell imaging and Pulse video analysis (Curi Bio)_55_ and Mantarray (Curi Bio)_57_ and Pulse video analysis (Curi Bio), respectively. **f,** Calcium transient recordings from WT and PKP2_Hom_ (n = 12 technical replicates each) assessed with CuriBio Nautilai (https://www.curibio.com/nautilai). Parameters displayed are: time to peak from 50% to peak (T_50_-Peak), Ca^2+^ transient width at 50% (CaD50), and the rate at which [Ca^2+]^_i_ reach its peak (Rise Rate). **g,** Acute PKP2 vs ACADVL silencing showed differential impact on field potential, contractility, and calcium transient of iPSC-CMs (details in Supplementary Fig. 4). Quantified data are presented as mean ± SD. Comparison p values were calculated by Prism unpaired t-test and ordinary One-Way ANOVA (Tukey’s post-hoc test) for **c** and **g**: p values: *p<0.05, **p<0.01, ***p<0.001, ****p<0.0001, ns = non-significant.

Perturbed electrophysiological properties of monolayers, determined by an extracellular recording of cardiac field potential using Maestro Pro Microelectrode array (MEA, Axion Biosystem), had been reported in our previous study^20^. Similarly, PKP2_Hom_ isogenic cells showed prolonged field potential duration and depressed spike amplitude and slope and depressed conduction velocity that were mostly correlated with copy numbers of mutated *PKP2* gene (Fig. 5d and Supplementary Fig. 3d). Contractile function of isogenic lines was characterized in either monolayer or 3D engineered heart tissue (EHT) platform. PKP2_Hom_ isogenic cells showed significantly impaired contractility than that of WT cells as quantified by the following parameters: contraction velocity, contraction amplitude, contraction and relaxation time for monolayers; and twitch force, contraction and relaxation velocity for 3D EHTs (Fig. 5e and Supplementary Fig. 3c). In addition, calcium transients were also disrupted in the PKP2_Hom_ isogenic line compared to the WT isogenic control, as shown by prolonged rise time from 50% to peak (t_50_-Peak) and Ca_2+_ transient width at 50% (CaD50), and slower rise rate (Fig. 5f).

Since iPSC-CMs used predominantly fatty acids for energy production as shown earlier (Fig. 4b and 4c), selective FAO disruption by ACADVL silencing altered electrophysiological properties (Fig. 5g and Supplementary Fig. 4a), contractility (Fig. 5g and Supplementary Fig. 4b), and Ca_2+_ transients (Fig. 5g and Supplementary Fig. 4c) that were all functionally connected with cellular structure, supporting the notion that energy metabolism is tightly integrated with structural functions^59^. So far, we demonstrated that both cKO heart tissue (Fig. 1, 2, and Supplementary Fig. 1, 2) and human PKP2 isogenic cardiomyocytes (Fig. 5a, b, and c) showed impaired FAO, glycolysis, and glucose oxidation. Functionally, we demonstrated how the impaired energy metabolism was manifested as depressed bioenergetics, consequently, supporting poor contractility in cardiomyocytes with compromised PKP2 function (Fig. 5d, e, and f). However, we had not confirmed whether reversing metabolic impairment can improve these structural functions that were compromised due to PKP2 mutation.

### Enhancing oxidative and glycolytic metabolism improved contractility but not Ca^2+^ transients of PKP2^Hom^ iPSC-CMs

The reversibility of metabolic impairment and the subsequent improvement of functional outcome was delineated by examining contractility, electrophysiological properties, and Ca_2+_ transients in PKP2_Hom_ isogenic cells (Fig. 6). As mentioned earlier, PKP2_Het_ cardiomyocytes, expressing some PKP2, could confound the data interpretation when examining functions such as contractility, calcium transient, and electrophysiological properties that have well understood structural connection to PKP2 and desmosome.

**Figure 6.**
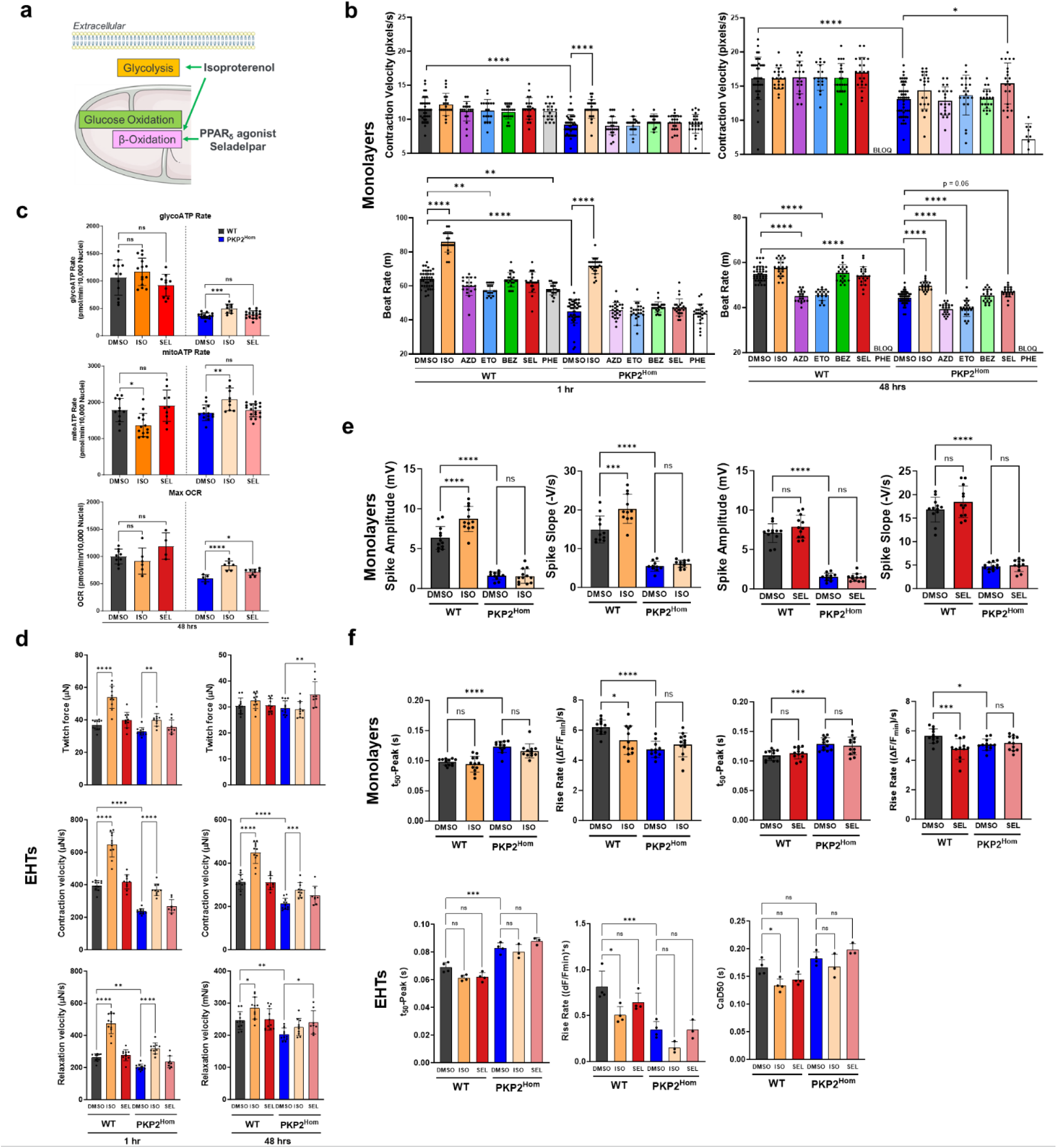
Enhancing oxidative and glycolytic metabolism improved contractility but not electrophysiological properties and Ca_2+_ transients of PKP2_Hom_ iPSC-CMs. **a,** Study design to evaluate small-molecule metabolic modulators in isogenic *Pkp2* homozygous cell model. Monolayers or engineered heart tissues (EHTs) were treated with isoproterenol or seladelpar for 1 and 48 hours before the measurement on contractility, bioenergetics, electrophysiological properties (field potential), and Ca_2+_ transients. **b,** WT and isogenic PKP2_Hom_ cells were treated pharmacologically with DMSO, isoproterenol (ISO), AZD7687 (AZD), etomoxir (ETO), bezafibrate (BEZ), seladelpar (SEL), and phenformin (PHE). Contractility was evaluated at 1 and 48 hrs post treatment using SONY live cell imaging and Pulse video analysis (n = 18-48 technical replicates) (Curi Bio)_55_. Average nuclear counts from live cells were used to normalize contraction velocity. **c,** In response to isoproterenol or seladelpar treatment, glycoATP and mitoATP rates and Max OCR of WT and isogenic PKP2_Hom_ cells were quantified using the Seahorse MitoStress test (n = 4-18 technical replicates). Statistical significance within the genotypes was determined at 48 hours post post-vehicle control/small molecule administration using Prism one-way ANOVA (Tukey’s post-hoc test). **d,** Assessment of normalized twitch force, contraction and relaxation velocity in WT and PKP2_Hom_ generated 3D-EHTs 1 hour or 48 hours post-vehicle control/small molecule administration (n = 8-11 technical replicates) using Mantarray (Curi Bio)_57_ and Pulse video analysis (Curi Bio)_55_. **e,** Assessment of field potential parameters, spike amplitude and spike slope, in WT and PKP2_Hom_ monolayers at 48-day post-vehicle control/small molecule administration using Maestro Pro Microelectrode array (MEA) platform (Axion Biosystems)_58_. Data were analyzed through AxIS Navigator software and the Cardiac Analysis Tool (Axion Biosystems) (n = 12 technical replicates). **f,** Calcium transient recordings from WT and PKP2_Hom_ monolayers (n = 12 technical replicates) or 3D engineered heart tissues (EHTs) (n = 3-4 technical replicates) were evaluated with CuriBio Nautilai (https://www.curibio.com/nautilai), multiwell format, and analyzed by Curi Bio’s Pulse platform_55_. Parameters shown are: time to peak from 50% to peak (T_50_-Peak), the rate at which [Ca^2+]^_i_ reach its peak (Rise Rate), and half-width/Ca_2+_ transient width at 50% (CaD50). Data are presented as mean ± standard deviation (s.d.). Statistical significance was determined using Prism one-way ANOVA (Tukey’s post-hoc test): p values are indicated as follows: *p<0.05, **p<0.01, ***p<0.001, ****p<0.0001.

We first tested a serial of metabolic enhancers or repressors, isoproterenol, AZD7687, etomoxir, bezafibrate, seladelpar, and phenformin, to examine their ability to modify contractility of PKP2_Hom_ isogenic cells (simplified Fig. 6a). As showed in Figure 1, mRNAs of the master regulators of cardiac energy metabolism, PPAR family members, were significantly changed in cKO heart tissue and restored by AAV9:PKP2. Since the three PPAR members each displayed distinct mRNA changes, we selected a pan-PPAR agonist, bezafibrate, used for treating metabolic syndrome with activities in reducing TAG levels and increasing both glucose and fatty acid metabolism^60^ as a tool to enhance overall energy metabolism in iPSC-CMs. Although PPARδ mRNA showed differential responses to the mode of AAV9:PKP2 treatment (Fig. 1d and Supplementary Fig. 1c) in *Pkp2-cKO* mouse heart, PPARδ activation is documented in its association with increasing total oxidative metabolism and glycolysis in cardiomyocytes^45^. We therefore selected a highly selective PPARδ agonist, seladelpar, to stimulate FAO and glycolysis and glucose oxidation^45^. Isoproterenol, a known chronotropic agent, served as a positive control for modulating bioenergetics^61^, contractile function^62^, electrophysiology^63^, and calcium transient^64^. Compared to isoproterenol treatment, seladelpar significantly improved contraction velocity and showed a trending improvement in beat rate of the PKP2_Hom_ monolayer at 48 hours of drug treatment, in contrast to no response to the pan-PPAR agonist bezafibrate (Fig. 6b). Compared to isoproterenol and seladelpar, AZD7687, etomoxir, or phenformin showed a general trend of suppressing beat rate in both WT and PKP2_Hom_ isogenic cells at 48 hours of drug treatment, suggesting metabolic modulation generates distinct functional outcome.

We observed a limited bioenergetic stimulation in WT or PKP2_Hom_ isogenic cells by isoproterenol or seladelpar using the standard Seahorse assays where the assay buffer contains BSA-palmitate as the only source for FAO. Statistical significance was reached when comparing bioenergetic responses within the genotypes at 48 hours of drug treatment. Isoproterenol stimulated basal glycoATP and mitoATP rates as well as Max OCR whereas seladelpar showed bioenergetic stimulation on Max OCR in PKP2_Hom_ isogenic cells (Fig. 6c). Similar to the improved contractility observed on the monolayer, we observed stimulated contractility in 3D EHTs of PKP2_Hom_ isogenic cells by either isoproterenol or Seladelpar as demonstrated by increased twitch force and contraction and relaxation velocity at either 1 hour or 48 hours of drug treatment (Fig. 6d). Seladelpar-mediated improvement on contractility was mainly observed on PKP2_Hom_ not WT isogenic cells. In contrast to expected response by isoproterenol treatment, seladelpar had little or no effect on electrophysiological properties of PKP2_Hom_ monolayers as characterized by spike amplitude and spike slope of field potential (Fig. 6e) and on calcium transient as characterized by t_50_-peak and rise rate of monolayers and by t_50_-peak, rise rate, and Ca^2+^ transient width at 50% (CaD50) of 3D EHTs (Fig. 6f).

Our studies showed that metabolic impairment due to PKP2 mutation partially responded to metabolic stimulation and subsequently resulted in improved contractility, whereas electrophysiological properties and calcium transients showed little or no response. These results revealed a new observation on differential sensitivity among structural functions of cardiomyocytes to metabolic perturbation. Therefore, energy-responsive contractile function can be asynchronized or decoupled from electrophysiological properties and calcium transients, likely presenting a higher risk, for example, when there is an increased metabolic demand by heart during exercise.

### TN-401 restored desmosome protein expression, contractility, electrophysiological properties, and Ca^2+^ transients in PKP2^Hom^ iPSC-CMs

Based on our results, we identified a mechanism leading to decoupling of contractility from electrophysiological properties and calcium transient in the presence of energy demand. Despite impaired PKP2 and desmosome function, contractile functions of cardiomyocytes remained able to respond and partially cooperated the need to match the energy demands.

To fully restore essential roles dictated by mechanical and structural integrity of PKP2 and desmosome, TN-401 was administered to PKP2_Hom_ isogenic cells, restoring PKP2 expression (Fig. 7a), as determined using immunofluorescence-based detection and quantification of PKP2 and DSP expression in a dose-dependent fashion (Fig. 7b). Contractile function of PKP2_Hom_ isogenic cells was rescued in a dose-dependent manner by TN-401, measured as displacement and velocity on monolayer (Fig. 7c). A single-dose rescue of contractility, measured by twitch force and contraction and relaxation velocity, was demonstrated with 3D EHTs (Fig. 7d). A single-dose rescue was achieved for electrophysiological properties measured by spike amplitude, spike slope, and conduction velocity; and for calcium transient measured by time to peak, calcium transient area under the curve, and calcium transient width at 50% (CaD50) (Fig. 7e). In addition, a single dose of AAV9:PKP2 renormalized the metabolite profiles and the integrated metabo-transcriptional network of cKO mouse heart (Fig. 1, 2, and Supplementary Fig. 1, 2).

**Figure 7.**
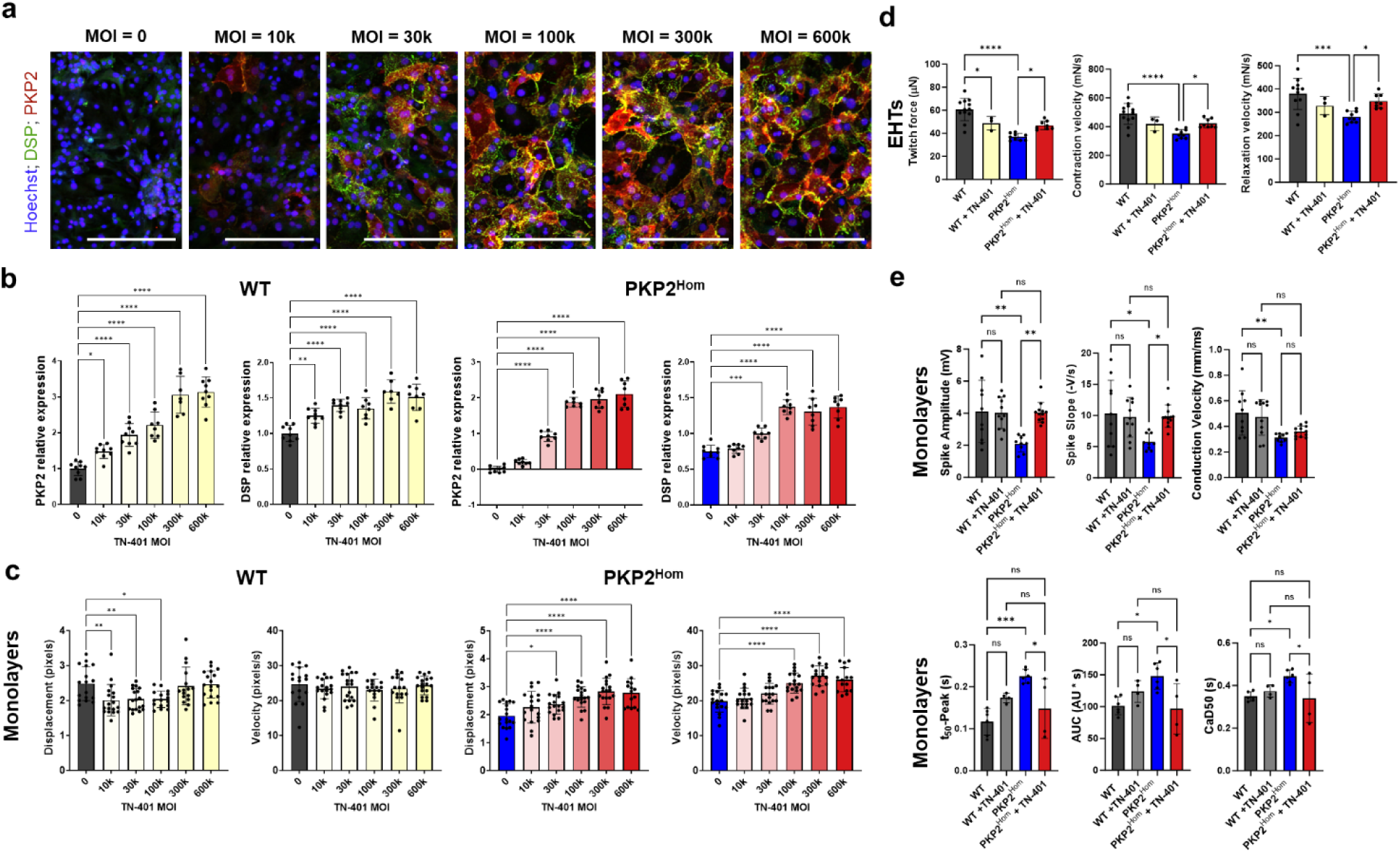
TN-401 treatment restored desmosome protein expression, contractility, electrophysiological properties, and Ca_2+_ transients in PKP2_Hom_ iPSC-CMs. **a**, Representative immunofluorescence images for PKP2 (red) and DSP (green) for the PKP2_Hom_ iPSC-CM isogenic line 21 days after transduction with TN-401 at the indicated MOIs. Nuclei were counterstained with Hoechst (blue). Scale bars = 200 mm. **b,** Quantification of PKP2 and DSP protein expression based on immunofluorescence staining for the WT and PKP2_Hom_ iPSC-CM isogenic lines. Protein expression was normalized to WT MOI 0 expression levels for both proteins; n = 7-9. **c,** Evaluation of contraction displacement and velocity in WT and PKP2_Hom_ iPSC-CMs at day 18 following administration of TN-401 at different MOIs: 0, 10000, 30000, 100000, 300000, and 600000 (n = 15-18) using SONY live cell imaging and Pulse video analysis (Curi Bio)_55_. **d,** Assessment of twitch force, contraction and relaxation velocity in WT and PKP2_Hom_ generated 3D-EHTs at day 15 (for WT) and 14 (for PKP2_Hom_) post-TN-401 administration, MOI = 100k; n = 12 and 3 for WT and WT + TN-401, respectively; n = 9 and 8, and PKP2_Hom_ and PKP2_Hom_ + TN-401, respectively, using Mantarray (Curi Bio)_57_ and Pulse video analysis (Curi Bio)_55_. **e,** Assessment of field potential parameters using Maestro Pro Microelectrode array (MEA) platform (Axion Biosystems)_58_ (top row, n = 9-12) and Ca_2+_ transient parameters (bottom row, n = 4-6) in WT and PKP2_Hom_ 11 days and 7 days post-TN-401 administration, respectively, using CuriBio Nautilai (https://www.curibio.com/nautilai), multiwell format, and analyzed by Curi Bio’s Pulse platform_55_. Monolayers were paced at 1.5 Hz during MEA recordings and at 1 Hz during Ca_2+_ transient measurement. Quantified data are presented as mean ± SD. Comparison p values were calculated by Prism one-way ANOVA (Tukey’s post-hoc test): p values: *p<0.05, **p<0.01, ***p<0.001, ****p<0.0001.

## Discussion

In this study, nontargeted metabolomics and lipidomics identified impaired metabolic pathways that responded to AAV9:PKP2-mediated restoration of PKP2 expression in *Pkp2-cKO* mouse heart tissue. Supported by metabolic dysregulation observed in human ARVC heart^18^, impaired lipid homeostasis, glycolysis, and glucose oxidation were confirmed in iPSC-CMs in response to either acute or stable disruption of PKP2 expression. Compromised PKP2 expression led to decreased contractile function and perturbed electrophysiological properties and Ca_2+_ transients in cardiomyocytes. We delineated the intrinsic cellular connection between metabolic impairment and cardiomyocyte dysfunction, showing that impaired energy production supported poor contractility, an effect that was exacerbated in PKP2 compromised cardiomyocytes. Furthermore, enhancing oxidative and glycolytic metabolism through pharmacological treatment with known metabolic enhancers improved contractile function but not electrophysiological properties and Ca_2+_ transients, indicating that essential roles dictated by mechanical and structural integrity of desmosome required restoration of PKP2 function as demonstrated by TN-401 gene replacement. Therefore, the data presented here support the interpretation that PKP2 serves to coordinate the energy metabolism of the cardiomyocyte to contractile function, while compromised PKP2 led to metabolic impairment and cardiomyocyte dysfunction at the intracellular level.

Our results demonstrated that structure-mediated functions, such as contractility, electrophysiological properties, and Ca_2+_ transients, differentially responded to metabolic perturbance. Therefore, metabolic impairment due to compromised PKP2 function may increase the risk of decoupling energy-responsive contractility from the less energy-responsive electrophysiological properties and Ca_2+_ transients, revealing a new arrhythmogenic mechanism that was intrinsically intracellular and in the course of disease likely initiated before the overt structural changes and/or occurring in the presence of energy challenges and metabolic stress. In the introduction section, we mentioned the exercise-triggered cardiac adversity in ARVC patients. Our data here showed that the loss of synchrony between energy-responsive contractility and the less energy-responsive electrophysiological properties and Ca_2+_ transients can be triggered by energy demand or metabolic stress at the cellular level, adding a new intracellular mechanism to the known arrhythmogenic triggers in ARVC, the disruption of gap junctions and ion channels and as the disease progresses, the epicardial fibrofatty replacement and/or scar tissue formation^65^.

### Metabolic impairment

In agreement with the published human ARVC tissue data^18^, we observed mRNA signatures of depressed FAO and glycolysis/glucose oxidation and increased fatty acid synthesis in cKO mouse heart (Fig. 1c and 1d). In addition, mRNA signatures of TAG dynamics were greatly perturbed with respect to fatty acid storage and mobilization via lipolysis. AAV9:PKP2 treatment showed significant long-term reversal of the metabolic impairment (Fig. 1c and 1d). In addition, reversal of metabolic impairment by AAV9:PKP2 demonstrated a dose-dependent response to PKP2 expression level, suggesting a direct connection between PKP2 function and the effects on cardiac energy metabolism (Supplemental Fig. 1).

We applied an established integrated omics approach, integrating nontargeted polar metabolomics and RNA sequencing within the same biological network^47,48^, to detect significant changes in metabolites and genes that coordinately responded to AAV9:PKP2 treatment. The integrated omics analysis identified metabo-transcriptional signatures that agreed with impaired FAO and glycolysis and glucose oxidation observed in human hearts with end-stage ARVC. Impaired TCA cycle was demonstrated by reduced intensities for ions annotated to citrate/isocitrate, malate and fumarate, and increased AKG in cKO mouse heart tissue (Fig. 2c). Reduced citrate in cKO heart might be due to consumption of that metabolite in the cytosol as a result of an increased *de novo* fatty acid synthesis, which was supported by upregulation of mRNA signatures in fatty acid synthesis (Fig. 1c). We hypothesized that increased fatty acid synthesis coordinated with reduced FAO could further exaggerate TAG content, consistent with our observation of increased lipid granules in iPSC-CMs in response to acutely or stably disrupted PKP2 expression, as well as increased TAG signals seen in lipidomics analysis of cKO mouse heart tissue (Fig. 1, Fig. 2, Fig. 3, Fig. 5a and b).

Intriguingly, mRNA of isocitrate dehydrogenase-2 (IDH2), catalyzing the conversion between isocitrate and AKG in the TCA cycle, was down-regulated, accompanied with decreased citrate/isocitrate signal and increased AKG in cKO mouse heart (Fig. 2c). AKG plays a key role in regulating amino acid synthesis, ATP production, redox homoeostasis, and metabolism-associated activities such as cellular signaling and epigenetic modifications^66,67^. As discussed in our previous published study^20^, restoration of a single component of the desmosome, PKP2, led to significant survival benefits of improved cardiac function, reversal of adverse RV remodeling, reduced severity and frequency of ventricular arrhythmia, and prevention of fibrosis. In this study, we demonstrated that “on-target” PKP2 effects extend beyond the desmosomes, at the transcriptional level, to energy metabolism. However, it remained unclear how disruption of PKP2 and desmosome functions signaled metabolic changes at the mRNA level. We postulate here that an early disruption of AKG homeostasis, a key component of the cells’ epigenetics machinery could possibly signal molecular changes to initiate a progressive metabolic impairment of ARVC through downstream changes in DNA methylation and associated longer-term changes in gene expression.

In response to acutely or stably disrupted PKP2 expression in iPSC-CMs, we confirmed impaired FAO and glycolysis and glucose oxidation by examining bioenergetics in the presence of known metabolic modulators. By delineating the intrinsic cellular connection between metabolic impairment and cardiomyocyte dysfunction, we showed that impaired energy metabolism led to suboptimal bioenergetics that supported poor contractility, which was exacerbated in PKP2 compromised cardiomyocytes. The metabolic impairment observed in both human and mouse ARVC heart tissue was recapitulated using isogenic lines carrying a known human pathogenic PKP2 mutation. Moreover, using PPARδ agonist, seladelpar, we observed some improvement in contractile function, mostly in the PKP2 homozygous cardiomyocytes but not in the WT isogenic cells. It is likely that this improvement in contractile function can be mediated by increasing fatty acid mobilization and oxidative and glycolytic metabolism despite the existing pathophysiological state; or by increasing metabolic flexibility to switch to other alternative fuels, such as medium-chain fatty acids or ketone bodies. Our studies suggested that ARVC cardiomyocytes may find it challenging to switch fuel source when both oxidative and glycolytic metabolism were simultaneously impaired. It remained unclear to us what energy fuel is opted for usage in ARVC hearts. Increased contribution of ketone bodies as an alternative fuel source has been shown in end-stage ARVC^18^, human failing hearts^25^, HCM^30^, and as an adaptive response to energy deprivation in animal models^68^. Song et al.^18^ reported upregulated ketone metabolic enzymes, OXCT1 and HMGCS2, suggesting increased ketone body synthesis at early-stage ARVC. Instead of increased ketone body usage, they reported that right ventricle at end-stage ARVC predominantly used medium-chain fatty acids as the mRNA encoding the medium-chain fatty acid oxidation enzyme, ACADM, was elevated in ARVC right ventricles. We observed down-regulated gene expression of *Oxct1*, *Hmgcs2*, and *Acadm* in our cKO mouse heart tissue^20^. We are currently investigating fuel flexibility of PKP2 heterozygous cell when medium-chain fatty acid and ketone bodies are provided as alternative fuel sources.

### Lipotoxicity

Our study agreed with early observations on increased intracellular lipid granules or lipid droplets detected in heart tissue of human or mouse model and in patient-specific ARVC iPSC-CMs^35–40^. Supported by clinical evidence, deranged cardiac lipid metabolism leads to intramyocardial increases of TAG and toxic lipids that have been linked to cardiac dysfunction in heart failure^26,29^. Increased oxidant production and DNA damage was reported in PKP2-deficient adult mouse heart and iPSC-CMs^16^.

Quantification of lipid peroxidation indicated higher oxidative stress in RV sections of ARVC patient hearts^38^. Our study focused on whether PKP2 coordinated the energy metabolism of the cardiomyocyte to contractile function and how compromised PKP2 led to metabolic impairment and cardiomyocyte dysfunction. We have not investigated whether impaired energy metabolism ultimately impacts mitochondria fitness and leads to progressive myocardial damage due to cardiomyocyte death and cardiac fibrosis.

Using a HFpEF mouse model, Yoshii et al.^69^ reported impact of reduced FAO on mitochondrial quality control, providing direct evidence of enhancing cardiac FAO to improve heart failure phenotypes. Stimulating energy metabolism to improve cardiomyocyte function was reported in homozygous iPSC-CMs carrying PLN-R14del where impaired FAO and preferred glycolysis led to enhanced lipid droplet storage.

Intriguingly, Harakalova et al.^70^ reported that activation of PPARα using bezafibrate improved mitochondrial structure and calcium handling function in iPSC-CMs harboring a PLN-R14del pathogenic mutation.

### Limitations of the study

There are several limitations inherent to this study. It is unclear how the PPARδ agonist seladepar improved contractile function and how metabolic modulation was communicated or mutually communicated with structural changes mediated by PKP2 and desmosome. Members of PPAR family play unique and overlapping roles in regulating energy homeostasis from individual cells to whole organism. Our study showed that PKP2 coordinated the energy metabolism of the cardiomyocyte to enable contractile function. Although impaired FAO and glycolysis and glucose oxidation have been reported in human ARVC heart tissue, we cannot anticipate how cardiac function would respond to metabolic stimulation. We have proposed a new intracellular arrhythmogenic mechanism and will need to conduct additional studies to estimate the associated arrhythmia risk when there is an increased metabolic demand by heart during exercise.

We acknowledged the metabolic immaturity and difference in Ca_2+_ handling of iPSC-CMs and challenges of conducting studies using adult rodent cardiomyocytes. To address this, we used Seahorse-based bioenergetic analysis to evaluate the glycolytic and oxidative state of iPSC-CMs and confirmed that fatty acid was the predominant energy fuel of iPSC-CMs in culture. In addition, 3D EHTs cast with isogenic cells were utilized to evaluate the contractile properties and Ca_2+_ transients in response to metabolic modulation.

As discussed in our early published work, the ARVC mouse model is a cardiac-specific conditional knock-out model of *Pkp2* with a severe phenotype. Therefore, the robust phenotype presented in this mouse model could possibly have different sensitivities to gene therapy relative to the more slowly developing phenotype associated with human disease. Our study modeled and revealed a direct intracellular connection between compromised PKP2 function and metabolic impairment that agreed with the human data. These findings require validation in patients with ARVC caused by *PKP2* mutations where gene therapy trials are ongoing.

## Conclusions

In this study, we utilized ARVC mouse and iPSC-CM models to confirm impaired lipid homeostasis, glycolysis, and glucose oxidation that concorded with the human data. We modeled and revealed a direct intracellular connection between compromised PKP2 function and metabolic impairment. Our studies indicated that structure-mediated functions, such as contractility, electrophysiological properties, and Ca_2+_ transients, differentially responded to metabolic perturbance. Increased risk of decoupling energy-responsive contractility from less energy-responsive electrophysiological properties and Ca_2+_ transient may be a new arrhythmogenic substrate occurring before the overt structural changes and/or in the presence of energy challenges. Reversing metabolic impairment by early intervention should improve cardiac energy metabolism and therefore, improve ARVC phenotypes. Metabolic biomarkers in plasma or cardiac tissue associated with the concealed phase of the disease may provide new screening tools to identify patients with increased risk of early adverse cardiac events. Our efficacy data with PKP2 gene therapy and reversal of metabolic impairment shed more light on understanding ARVC etiology and in practice, may help patients and physicians to make decisions regarding disease management and treatment.

## Acknowledgement

We thank Dr. Mario Delmar and NYU for licensing *Pkp2-cKO* mouse.

## Contributions

Conceptualization: Z.J.Y. and J.A.A.; Methodology: J.A.A., A.E.T., I.P.-G., D.H.-S., B.R., K.F., E.D., K.W., I.W., A.Z., M.VP., E.X., J.W., S.J., Y.H., C.D.-H., J.M., K.R., and Z.J.Y.; Formal analysis: J.A.A., A.E.T., I.P.-G., D.H.-S., K.F., E.D., K.W., I.W., and Z.J.Y.; RNA sequencing analysis: R.S. and F.F.; Multiomics and lipidomics analyses: D.H.-S., K.F., E.D., K.W., and J.A.A.; Investigation: J.A.A., A.E.T., I.P.-G., D.H.-S., E.D., Z.J.Y.; Data curation: J.A.A., A.E.T., I.P.-G., D.H.-S., B.R., K.F., E.D., K.W., and Z.J.Y.; Writing original draft preparation: Z.J.Y.; Writing, Review, and Editing: Z.J.Y., E.D., J.A.A., A.E.T., I.P.-G., D.H.-S., J.P., A.G.-S., T.H.; Supervision: Z.J.Y., E.D., G.A., A.G.-S., J.P., F.F., S.J. X.S., K.N.I., and T.H..

