## Supplemental materials and methods for "Plakophilin-2 Coordinates Energy Metabolism and Contractility in Cardiomyocytes, Revealing Its Roles beyond Desmosomes"

### Supplementary Materials

**Supplementary Figure 1. AAV9:PKP2 gene therapy largely maintained key cardiac energy metabolism that was perturbed in *Pkp2-cKO* ARVC mouse model**

**Supplementary Figure 1. AAV9:PKP2 gene therapy largely maintained key mRNA signatures of cardiac energy metabolism that was perturbed in *Pkp2-cKO* ARVC mouse model**

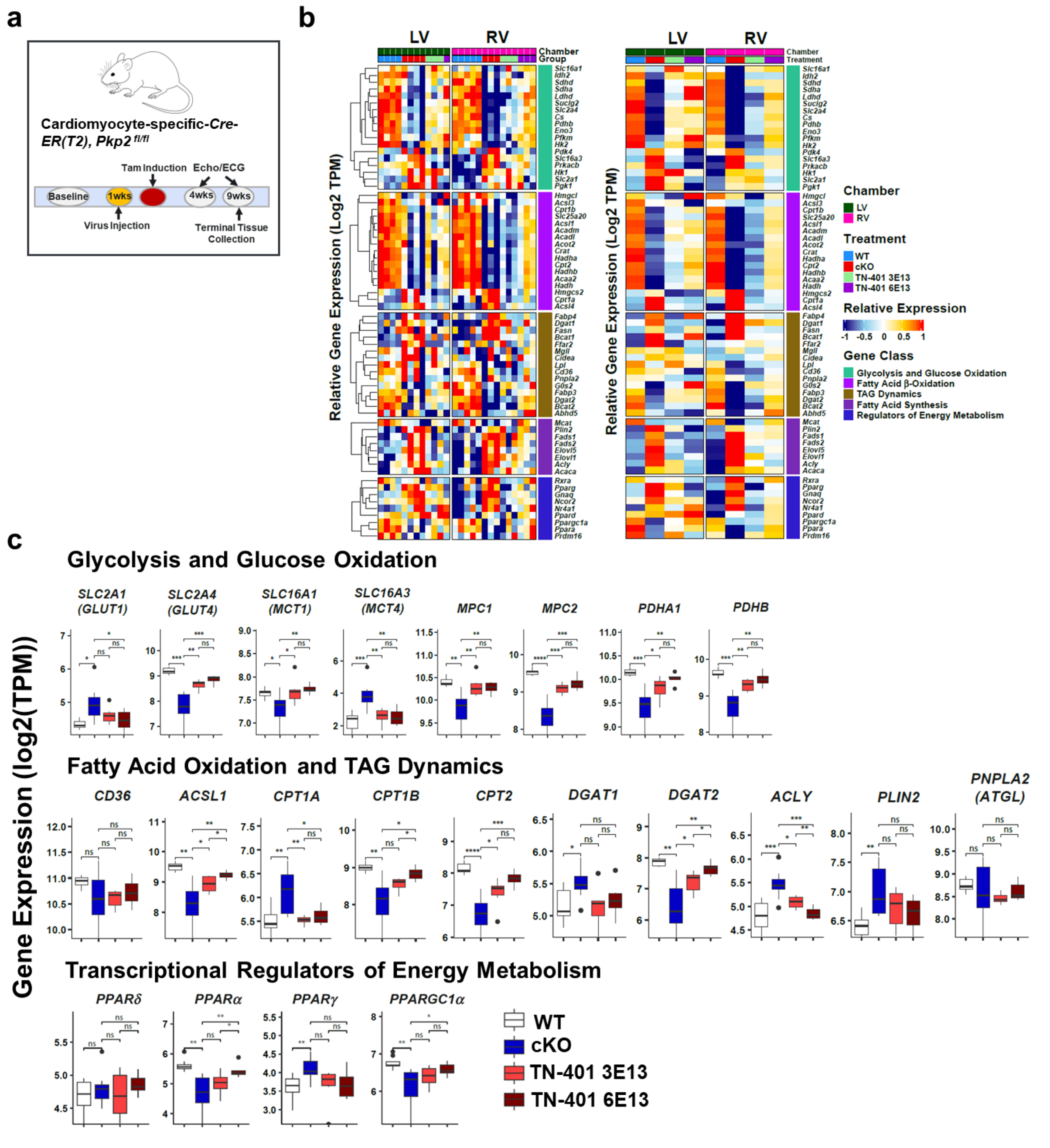

**a,** Study design to evaluate TN-401 dose-dependent efficacy at week 4 and 9 post tamoxifen induction of *Pkp2* gene deletion in *Pkp2-cKO* mouse model. Animals were dosed by TN-401 at 3E13 or 6E13 vg/kg at 1 week before induction and heart tissue collected at week 10 post virus injection. **b,** Relative gene expression of selected genes was measured by RNA-seq. Heatmap of gene expression was sorted by heart chambers (LV vs RV), treatment groups, and gene classes. Left panel shows heatmap of gene expression of each individual animal and its corresponding chambers presented in scaled log2-transformed. Right panel shows a scaled average across samples of each treatment group. Numbers of samples in each treatment group used for RNA sequencing (for subsequent integrated omics) were 9 WT, 7 cKO, 6 and 4 cKO+TN-401 at 3E13 and 6E13 vg/kg, respectively, including both RV and LV. **c,** Boxplots showed group-wise gene expression for of the selected gene classes. Each box showed the distribution of expression values in the following manner: the midline represented the median expression value, the box indicated the interquartile range where the middle 50% of values lie, and the whiskers at the top and bottom of each box represented the range of values outside the interquartile range. The black dots represent values that fall outside the 2$^{nd}$ and 3$^{rd}$ quartiles. Values were log 2 of TPM (Transcripts Per Million) and were aggregated from LV and RV. Comparison p values were calculated by Student's t-test: p values: *$p<0.05$, **$p<0.01$, ***$p<0.001$, ****$p<0.0001$.

**Supplementary Figure 2. Integrated metabolomics and transcriptomics showed AAV9:PKP2 maintained a broad spectrum of metabolites and their corresponding transcriptional signatures that were disturbed in *Pkp2-cKO* cardiac tissue**

**Supplementary Figure 2. Integrated metabolomics and transcriptomics showed TN-401 maintained a broad spectrum of metabolites and their corresponding transcriptional signatures that were disturbed in *Pkp2-cKO* cardiac tissue**

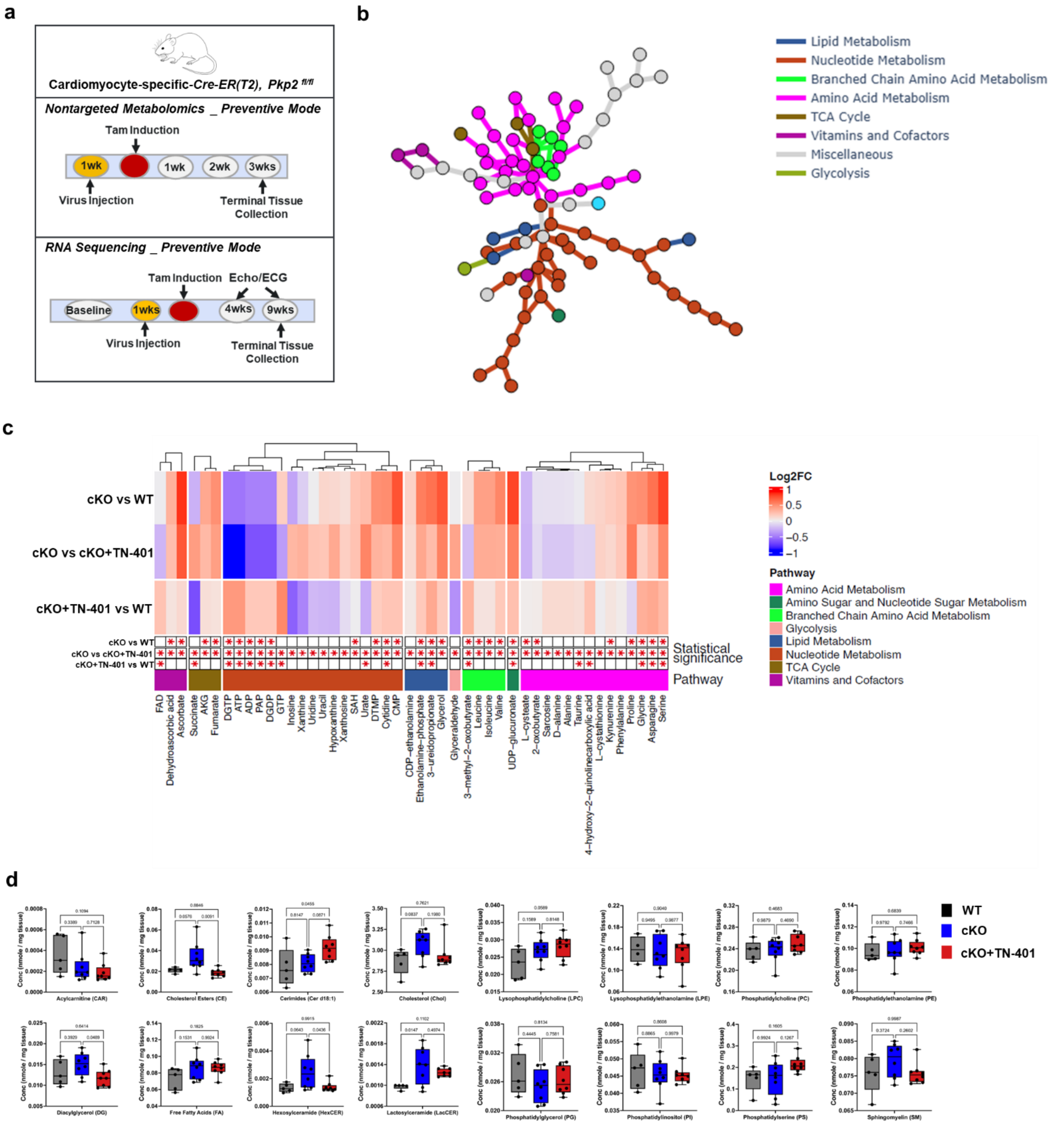

**a,** Study designs for nontargeted metabolomics and RNA sequencing, respectively, to evaluate AAV9:PKP2 efficacy in *Pkp2-cKO* ARVC mouse model. For nontargeted metabolomics, TN-401 was dosed at 1E14 vg/kg at 1week before induction (the preventive mode of treatment). RNA sequencing analysis used the dataset of the preventive mode as shown in Supplementary Figure 1. **b,** Integrated metabolomics and RNA sequencing showed that TN-401 in the preventive mode significantly maintained multiple metabolic pathways with steady-state metabolites and mRNA detected when compared to the untreated *Pkp2-cKO* cardiac tissue. **c,** The heatmap showed mean values of log2 fold change of metabolites with each column representing cKO vs WT, cKO vs TN-401 treated, and TN-401 treated vs WT, respectively. Metabolites were categorized by functional pathways. **d,** Average of additional lipid subclasses in cardiac tissue of WT, cKO, and TN-401 treated. Black dots represent the class average from each animal, while the midline represents the median of all animals, and the box indicates the interquartile range where the middle 50% of values lie, and the whiskers at the top and bottom of each box represent the range of values outside the interquartile range. Numbers of samples in each treatment group used for RNA sequencing and integrated omics were 9 WT, 7 cKO, 4 cKO+TN-401 at 6E13 vg/kg, respectively, including both RV and LV (the preventive mode). Number of animals in each treatment group used for metabolomics and integrated omics were 5 WT, 8 cKO, 8 cKO+TN-401 at 1E14 vg/kg (the preventive mode).

**Supplementary Figure 3. Characterization of isogenic human iPSC-CMs carrying heterozygous and homozygous pathogenic PKP2 mutation**

### Supplementary Figure 3. Characterization of isogenic human iPSC-CMs carrying heterozygous and homozygous pathogenic PKP2 mutation

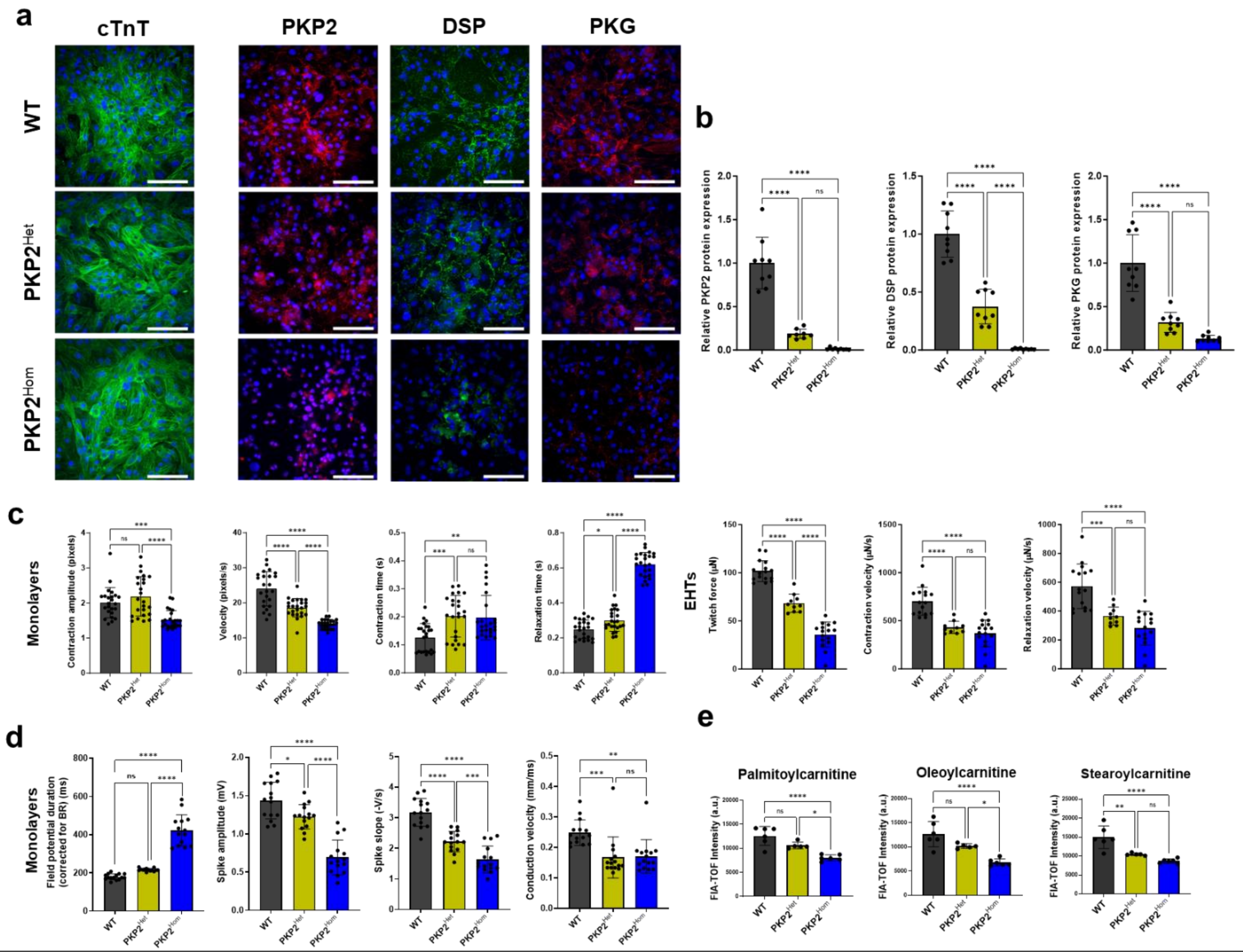


**a,** Detection of cTnT, PKP2, DSP, and PKG expression by immunofluorescence on mutant isogenic human iPSC cardiomyocytes carrying a pathogenic mutation c.2146G>C: PKP2 heterozygous mutant ($PKP2^{Het}$, first row), and PKP2 homozygous mutant ($PKP2^{Hom}$, second row); for the cardiac troponin T (first column), PKP2 (second column), DSP (third column), and PKG (fourth column). Scale bars = 100 mm. **b,** Quantification of PKP2, DSP, and PKG protein expression for the WT and the two mutant isogenic lines. Results were normalized to the WT protein expression levels; n = 8 technical replicates for $PKP2^{Het}$; n = 9 technical replicates for WT and $PKP2^{Hom}$. **c,** Monolayers (n = 24) and engineered heart tissues (EHTs) (n = 10 technical replicates for $PKP2^{Het}$, n = 16 technical replicates for $PKP2^{Hom}$) showed reduced contraction force and velocity, and prolonged contraction and relaxation time in the $PKP2^{Hom}$ lines using SONY live cell imaging and Pulse video analysis (Curi Bio)[55] and Mantarray (Curi Bio)[57] and Pulse video analysis (Curi Bio)[55], respectively. **d,** Monolayers MEA parameter characterized by prolonged field potential duration (FPD) along with depressed spike amplitude, conduction velocity, and spike slope using the Maestro Pro Microelectrode array (MEA) platform (Axion Biosystems)[58], n = 15 technical replicates. Quantified data are presented as mean ± SD. **e,** FIA-TOF mass spectrometry measurement showed reduced acylcarnitines in $PKP2^{Hom}$ compared to $PKP2^{Het}$ (n = 6 technical replicates, 3 wells of 12-well plate were pooled as 1 technical replicate). Comparison p values were calculated by Prism ordinary One-Way ANOVA (Tukey's post-hoc test): p values: *p<0.05, **p<0.01, ***p<0.001, ****p<0.0001, ns = non-significant.

**Supplementary Figure 4. Acute PKP2 vs ACADVL silencing showed differential impact on contractility, field potential, and calcium transients of iPSC-CMs**

### Supplementary Figure 4. Acute PKP2 vs acute ACADVL silencing showed differential impact on electrophysiological properties, contractility, and calcium transients of iPSC-CMs

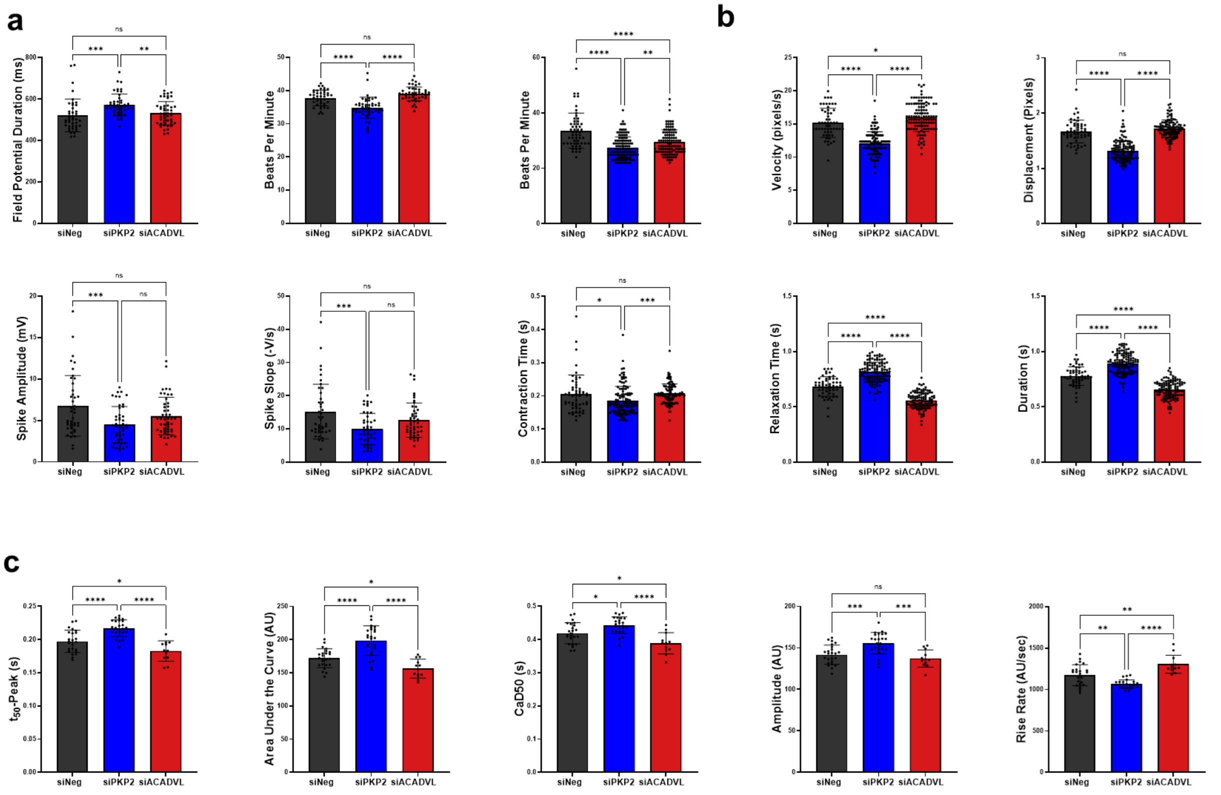


**a,** Cardiac field potential recordings were obtained from cardiomyocyte monolayers 7 days post siRNA silencing using the Maestro Pro Microelectrode array (MEA) platform (Axion Biosystems) [57]. Raw data were analyzed through AxIS Navigator software and the Cardiac Analysis Tool (Axion Biosystems) (n = 48 technical replicates). **b,** Contractility was evaluated at day 7 post siRNA silencing using SONY live cell imaging and Pulse video analysis (n = 54-111 technical replicates) (Curi Bio)[55]. Average nuclear counts from live cells were used to normalize contraction velocity. **c,** Cardiac calcium transients were evaluated in a multiwell mapping format using CuriBio Nautilai (https://www.curibio.com/nautilai). Raw data were analyzed by Curi Bio's Pulse platform (n = 12- 24 technical replicates). Quantified data are presented as mean ± SD. Statistical significance was determined using one-way ANOVA (Tukey's post-hoc test), performed in Prism: p values are indicated as follows: *p<0.05, **p<0.01, ***p<0.001, ****p<0.0001.

### Supplementary Materials

### Methods

#### Animal studies

Animal studies were performed according to Tenaya Therapeutics' animal use guidelines. The animal protocols were approved by the Institutional Animal Care and Use Committee (IACUC number: 2020.007).

Animals were allowed to acclimate for at least 3 days following shipment. Animals were housed in Innovive racks and disposable cages. The cages are pre-filled with alpha-dri as bedding. Our enrichments have nestlets and Twist-n'Rich for single-housed mice.

Before animal studies were initiated, study designs were created and reviewed that included the objective/hypothesis, treatment dose, sample size, primary outcomes, time points, etc. Baseline (before induction) body weight, age, and gender were used for inclusion and exclusion criteria. Study cages were kept on the same rack throughout the duration of the studies to minimize potential confounders.

For animal studies, ejection fraction was the primary outcome measurement used to determine sample size. In addition, the animal sample size per treatment group was determined based on our in-house experience of conducting efficacy, dose-ranging, and long-term survival benefit studies using AAV9:transgene as test article.

Animals were monitored daily. Humane endpoint was determined based on a body condition score, with weight loss, posture, respiration, activity, and mentation as factors. Animal death was documented and plotted in Kaplan-Meier survival curves and body weight curves. Animals that were euthanized or died prior to the last echocardiography or electrocardiogram (ECG) were excluded from the time progression curve of %

ejection fraction, RV size, and arrhythmia scores. Animals found dead were excluded from tissue DNA, RNA, or protein analyses. If an animal was found dead right after echo or ECG, it would be included in terminal tissue processes.

Standard operating procedure (SOP) was strictly followed. The reproducibility was ensured by using animals that were age-matched with evenly distributed body weight and sex. Tamoxifen induction and AAV9 injection were performed by one scientist and echocardiography and ECG were performed by another scientist for consistency. ECG raw traces were examined and double checked by two different scientists. AAV9 was produced and purified following the SOP and each lot titer was determined based on vector genomes. All attempts on replication were successful.

Animals were randomized for either the vehicle control or the AAV9-treated based on body weight and sex that were evenly distributed among treatment groups.

All *in vivo* animal studies were conducted blinded. Scientists were blinded to group allocation during data collection and analysis.

### Mouse model and route of AAV administration

Tenaya licensed a cardiomyocyte-specific, tamoxifen-activated *Pkp2-cKO (αMyHC-Cre-ER(T2)/Pkp2$^{fl/fl}$)* mouse line in the C57BL/6 background from Dr. Mario Delmar, NYU Grossman School of Medicine[1]. The *Pkp2-cKO* animals were induced with tamoxifen at 0.1mg/g for 3 consecutive days. Tamoxifen injection activates Cre recombinase in Cre-positive animals and induces homozygous deletion of *Pkp2* gene. TN-401 or AAV9:mPkp2 was given as a single dose via retro-orbital injection before or after animals were induced with tamoxifen. *Pkp2$^{fl/fl}$ Cre*-negative littermates were used as wild-type controls. HBSS, as vehicle control, was injected to WT or non-AAV treated *Pkp2-cKO* animals.

For Figure 1, WT or *Pkp2-cKO* animals aged 6-8 months were randomized for either the vehicle control or the AAV9-treated based on body weight and gender that were evenly distributed among treatment groups: (WT, HBSS: 4M/5F) (cKO, HBSS: 4M/5F) (cKO+AAV9:mPkp2 at 3E13 vg/kg, injection 1 week after KO induction: 5M/5F) (cKO+AAV9:mPkp2 at 1E14 vg/kg, injection 1 week after KO induction: 5M/5F) (cKO+AAV9:mPkp2 at 1E14 vg/kg, injection 2.5 week after KO induction: 6M/5F) with animal takedown at 50 weeks after virus injection.

For Supplementary Figure 1, WT or *Pkp2-cKO* animals aged about 6 months: (WT, HBSS: 3M/3F) (cKO, HBSS: 4M/4F) (cKO+TN-401 at 3E13 vg/kg, injection 1 week before KO induction: 4M/5F) (cKO+TN-401 at 6E13 vg/kg, injection 1 week before KO induction: 3M/3F) with animal takedown at 10 weeks after virus injection. Numbers of samples in each treatment group used for RNA sequencing and integrated omics were 9 WT, 7 cKO, 6 and 4 cKO+TN-401 at 3E13 vg/kg and 6E13 vg/kg, respectively, including both RV and LV.

For Figure 2, numbers of animals used for RNA sequencing were described in Figure 1. For nontargeted polar metabolomics of the therapeutic mode of treatment, WT or *Pkp2-cKO* animals aged about 4.5 months: (WT, HBSS: 5M/4F) (cKO, HBSS: 6M/2F) (cKO+TN-401 at 1E14 vg/kg, injection 2.5 week after KO induction: 6M/2F) with animal takedown at 2.5 weeks after virus injection. For shotgun lipidomics of the preventive mode of treatment, WT or *Pkp2-cKO* animals aged about 4.5 months: (WT, HBSS: 2M/3F) (cKO, HBSS: 5M/3F) (cKO+TN-401 at 1E14 vg/kg, injection 1 week before KO induction: 5M/3F) with animal takedown at 4 weeks after virus injection.

For Supplementary Figure 2, numbers of samples used for RNA sequencing were described in Supplementary Figure 1 above. For nontargeted polar metabolomics of the preventive mode of treatment, WT or *Pkp2-cKO* animals aged about 4.5 months: (WT, HBSS: 2M/3F) (cKO, HBSS: 5M/3F) (cKO+TN-401 at 1E14 vg/kg, injection 1 week before KO induction: 5M/3F) with animal takedown at 4 weeks after virus injection.

### AAV virus production

AAV production was carried out as previously described[2]. Briefly, HEK293T cells were seeded in a HyperFlask (Corning) and triple transfected using a 2:1 PEI:DNA ratio (PEI Max) with a helper plasmid containing adenoviral elements (pHelper), a plasmid containing the Rep2 and Cap genes from the respective AAV serotype, and finally an ITR containing plasmid to be packaged. Three days following transfection cells were harvested and lysed. Virus was purified using iodixanol ultracentrifugation and cleaned and concentrated in Hank’s Balanced Salt Solution (HBSS) + 0.014% Tween-20 using a 100-kDa centrifuge column (Amicon). AAV was titered using either a PicoGreen or ddPCR assay. AAV particles were diluted to proper titers on the day of animal injection in a formulation buffer, HBSS plus 0.001% Pluronic F68 as a surfactant. The formulation buffer is the carrier buffer for TN-401 or AAV9:mPkp2 to prevent aggregation of capsids. It serves as the vehicle control for the animals ‘without treatment’ as compared to animals ‘with AAV treatment’.

### Untargeted polar metabolome profiling

Polar metabolites were extracted from heart tissue and human iPSC-CM culture as described below:

Tissue:

Heart tissue was immediately snap frozen in liquid nitrogen upon removal. Metabolite extraction was performed on 30 – 80 mg frozen samples. Frozen samples with metallic beads were homogenized using the Storm Bullet Blender (Next Advance) in 0.5 ml of 70% ethanol (v/v in water) pre-chilled at ≤ -20°C. After homogenization, samples were transferred to pre-chilled 15 ml tubes, kept in cold ethanol bath, and followed by a hot extraction by mixing with 7 ml of 70% ethanol at 75°C. After 1 minute, the tube was vortexed quickly and then moved to cold ethanol bath. The 15 ml tubes were centrifuged for 10 minutes at 4000 rpm at ≤ 4°C. The supernatants were then transferred, stored at -80°C, and were analyzed within 30 days post-extraction. Samples from the same study were kept frozen and processed together on the same day.

Human iPSC-CM:

Cultured human iPSC-CMs were seeded at about 125,000 cells per well. Three wells of 12-well plates were pooled to form 1 technical replicate and a total of 6 such technical replicates per treatment, siNeg or siPKP2, and per genotype, WT, $PKP2^{Het}$, or $PKP2^{Hom}$, were harvested at approximately 70% confluence. Approximate cell number count was conducted using a sister plate before freezing all samples. Solvent extraction, using 40:40:20 proportion of acetonitrile: methanol: water, from all cell culture plates were done on the same day, identical timing, and the same extraction and wash solvent preparation.  Extracts were than kept at -80ºC until analysis.  Extracts were analyzed within 30 days post-extraction.

Metabolome profiles of the sample extracts are acquired using flow-injection mass spectrometry.  The method is adapted from Fuhrer[3].  The instrumentation consists of an Agilent 6550 iFunnel LC-MS Q-TOF mass spectrometer in tandem with an MPS3 autosampler (Gerstel) and an Agilent 1260 Infinity II quaternary pump.  The running buffer is a solution of 60:40 isopropanol in water (v/v) (OptimaTM LC/MS grade) buffered with 1 mM ammonium fluoride (Sigma-Aldrich).  Hexakis (1H, 1H, 3H-tetrafluoropropoxy)-phosphazine) (Agilent HP) and 3-amino-1-propanesulfonic acid (HOT) (Sigma-Aldrich) are added to the running buffer to serve as lockmasses.  The isocratic flow rate is 0.150 mL/min. The instrument is run in 4GHz High Resolution, negative ionization mode.  Mass spectra between 50 and 1,000 m/z are collected in profile mode.  5 μL of each sample are injected twice, consecutively, within 0.96 minutes to serve as technical replicates.  The pSS was injected periodically throughout the batch to serve as quality control.  Samples are acquired randomly within 96-well plates.

Raw profile data are centroided, merged, and recalibrated using methods adapted from Fuhrer[3].  Putative annotations are generated based on compounds contained in the Human Metabolome Database, KEGG, and ChEBI databases using both accurate mass per charge (tolerance 0.001 m/z) and isotopic correlation patterns.

### Network-based integration of metabolomics and transcriptomics data

Differential analysis data for metabolomics and transcriptomics data (see above) were subjected to multiomics analysis using the previously published R framework[4].  Briefly, a proprietary chemical reaction network was used to map annotated metabolites and transcripts onto the metabolic network.  The method’s algorithm then identified the most regulated metabolic subnetwork of an appreciable size as controlled by false discovery rate parameters.  A regulated subnetwork refers to a contiguous portion of the graph representation of metabolism where significant differences in both metabolite and transcript levels are seen between comparison groups. A parameter scanning procedure was used to identify values for missing metabolite and transcript penalties which generated the most consistent network sizes. Those regulated subnetworks chosen based on that parameterization were then represented with nodes representing metabolites and edges representing enzymes. In addition to a visualization where the log2 transformed fold-changes and p-values were represented, network representations highlighting metabolic pathway membership were also generated. For comparisons where few metabolites or transcript changes were observed in any part of the metabolic network, no regulated subnetwork was found, and those comparisons were reported as having no identifiable regulated network.

### Visualization of Metabolites Changes from Integrated Analysis

To illustrate the relative changes in metabolites identified through integrated metabolomics and transcriptomics analysis, we generated a heatmap using the *ComplexHeatmap* package in *R*. The heatmap depicts $log_2$-transformed fold changes and associated p-values for metabolites that were significantly altered in the TN-401 treatment group compared to the cardiac KO group. Metabolites were grouped according to their pathway and class annotations for clearer interpretation, and were clustered in each group using Euclidean distance and complete clustering method. Statistically significant differences within each comparison are marked with asterisks. The heatmap for the therapeutic mode of treatment is presented in Figure 2, while the heatmap for the preventive mode of treatment is shown in Supplementary Figure 2.

### Shotgun Lipidomics

For homogenized tissue, 20-100 mg of tissue were collected in a 2mL homogenizer tube pre-loaded with 2.8 mm ceramic beads (Omni #19-628). Then 0.75mL PBS was added to the tube and homogenized in the Omni Bead Ruptor Elite (3 cycles of 10 seconds at 5 m/s with a 10 second dwell time). Homogenate containing 1-8mg of original tissue was transferred to a glass tube for extraction. A modified Bligh and Dyer extraction[5] was carried out on all samples. Prior to biphasic extraction, a standard mixture of 75 lipid standards based on Avanti Ultimate Splash ONE was used (Avanti

330820, 861809, 330729, 330727, 791642). Following two successive extractions, pooled organic layers were dried down in a Thermo SpeedVac SPD300DDA using ramp setting 4 at 35 degrees C for 45 minutes with a total run time of 90 minutes. Lipid samples were resuspended in 1:1 methanol/dichloromethane with 10 mM Ammonium Acetate and transferred to robovials (Thermo 10800107) for analysis.

Samples were analyzed on the Sciex 5500 with DMS device (Lipidyzer Platform) with an expanded targeted acquisition list consisting of 1450 lipid species across 17 subclasses. Differential Mobility Device on Lipidyzer was tuned with EquiSPLASH LIPIDOMIX (Avanti 330731). Data analysis performed on an in-house data analysis platform comparable to the Lipidyzer Workflow Manager[6]. Instrument method including settings, tuning protocol, and MRM list available[6]. Quantitative values were normalized to mg of tissue.

### Genome Editing in human iPSC

CRISPR Cas9 mediated knock-in to introduce the pathogenic missense mutation c.2146-1G>C in *PKP2* gene in a human healthy iPSC line was generated by EditCo Bio, Inc. To generate these cells, ribonucleoproteins containing the Cas9 protein and synthetic chemically modified guide RNA produced by Synthego (guide RNA sequence: AAAACAGAUGCCGACAUCAG) were electroporated into the cells using EditCo's optimized protocol. Editing efficiency is assessed upon recovery, 48 hours post electroporation. Genomic DNA is extracted from a portion of the cells, PCR amplified (Forward primer: TCACCAATTTCATTCTGCAGAGA; Reverse primer: AGACTAGTTCCATCACAGTTCTAAT) and sequenced using Sanger sequencing using the reverse primer. The resulting chromatograms are processed using EditCo's Inference of CRISPR edits software (ice.synthego.com). To create monoclonal cell populations, edited cell pools are seeded at 1 cell/well using a single cell printer into 96 or 384 well plates. All wells are imaged every 3 days to ensure expansion from a single-cell clone. Clonal populations are screened and identified using the PCR-Sanger-ICE genotyping strategy described above.

### Human iPSC culture and differentiation to cardiomyocytes

Human iPSC isogenic lines were thawed and recovered in 6-well cell culture plate (Corning) according to manufacturer's instructions (Synthego). Human iPSCs were plated on Matrigel-coated plates (Corning) and expanded using mTeSR Plus medium (StemCell Technologies) under standard incubation conditions (37°C, 5% $CO_2$), with medium change every other day. When cell culture reached an 80-90% confluence, human iPSCs were dissociated with Dulbecco's Phosphate Buffered Saline (DPBS) solution (Gibco) supplemented with 0.5 mM UltraPure EDTA (Invitrogen) and passaged onto new Matrigel-coated 6-well plates for further expansion, or onto Matrigel-coated 632 $cm^2$ Nunc Cell Factories (Thermo Fisher Scientific) for differentiation to cardiomyocytes, in mTeSR Plus medium with 10 μM Y-27632 (BioOrbyt).

Differentiation of human iPSCs towards cardiomyocytes was conducted following a small molecule Wnt-activation/inhibition protocol [7]. Medium was changed to RPMI supplemented with B27 without insulin (Gibco) and 7 μM CHIR-99021 (LC Laboratories). After 48 hours, medium was replaced by RPMI with B27 without insulin for one day, then switched to RPMI supplemented with B27 without insulin and 5 μM IWP-2 (BioOrbyt). 48 hours following the addition of IWP-2, medium was changed back to RPMI supplemented with B27 without insulin for 48 hours, then replaced by RPMI supplemented with B27 with insulin (Gibco). 48 hours later, and upon observation of beating cardiomyocytes, selection of this cell population was carried out with Blasticidin (the three human iPSC isogenic lines exhibit Blasticidin resistance regulated by the cardiac myosin heavy chain 6 (MYH6) promoter), switching medium to RPMI supplemented with B27 with insulin and 2 μg/mL Blasticidin (Thermo Fisher Scientific), refreshing every 48 hours afterwards. On day 14, cardiomyocytes were dissociated using TrypLE Express (Gibco) and replated in RPMI supplemented with B27 with insulin and 10% fetal bovine serum (Gibco) onto new Nunc Cell Factories on a 2:1 ratio. 24 hours post-seeding, medium was switched back to RPMI plus B27 with insulin and 2 mg/mL Blasticidin, and refreshed every 48 hours. On day 21, cardiomyocytes were frozen in CryoStor CS10 cryopreservation medium (StemCell Technologies) at a 10 million cell/mL concentration, and stored in liquid nitrogen until further use.

### Engineered Heart Tissues (EHTs) casting and culture

EHTs were prepared using the Mantarray kit (CuriBio) and following manufacturer's instructions. Briefly, posts were pretreated with a 0.1% polyethylenimine (Sigma) solution for 10 minutes, washed with distilled water (Gibco), and followed by a second pretreatment with a 0.01% glutaraldehyde (Sigma) solution for 30 minutes. After two more distilled water washes, the Mantarray kit was reassembled and stored at 4$^{o}$C until EHTs were cast. On casting day, the casting plate was removed from 4$^{o}$C and placed onto a cold block. A 50 mL solution containing 3 mL of thrombin (stock concentration 100 U/mL, Sigma) and 47 mL of EHT medium (RPMI medium supplemented with B27 with insulin and 5 g/L 6-aminocaproic acid (Sigma)) was pipetted per well and returned to 4$^{o}$C until casting. A 100 mL mix containing human iPSC-CM (60 mL of a 8.3 x $10^6$ cells/mL suspension in EHT medium with 10 μM Y-27632), human cardiac fibroblasts (30 mL of a 2.5 x $10^6$ cells/mL suspension in EHT medium with 10 μM Y-27632) and 10 mL fibrinogen (stock concentration 50 mg/mL, Sigma) was prepared per well, then the Mantarray kit was removed from 4$^{o}$C and placed over a cold block. 100 mL of the previously prepared cell suspension were added per well and gently mixed with the 50 mL thrombin solution. The Mantarray kit with the seeded EHTs was incubated (37$^{o}$C, 5% $CO_2$) for 80 min, followed by the addition of 1 mL of EHT medium and another incubation for 10 min. After the second incubation was completed, the EHTs were transferred to a fresh 24-well plate (Greiner) with 2 mL EHT medium with 10 μM Y-27632 per well. For the first week, EHT medium was refreshed (2 mL/well) every other day. On day 8, medium was switched to metabolic maturation medium[8] and EHTs were maintained in this medium for at least 14 days, refreshing every 48 hours.

### iCell Cardiomyocytes$^{2}$ culture

iCell Cardiomyocytes$^{2}$, CDI CM$^{2}$, were thawed according to the manufacturer's instructions (FUJIFILM Cellular Dynamics) and seeded onto Matrigel-coated plates at a density of 20,000 cells per well of 96-well plates or at a seeding density proportional to the well size. Seeded cells were maintained in CDI maintenance media (FUJIFILM Cellular Dynamics) for 7-10 days until the day of treatments.

### siRNA knockdown of iPSC-CMs

Endogenous *PKP2* expression was knocked down based on the protocol described in Wu[9]. To knock down *ACADVL* expression in CDI CM[2], cell monolayer in CDI maintenance media was transfected with 1.25nM siRNA targeting *ACADVL* (Ambion by Life Technologies, Silencer™ Select pre-designed and validated) or Silencer™ Select negative control siRNA using Lipofectamine RNAiMAX transfection reagent (Invitrogen) mixed with Opti-MEM (Gibco), following the manufacturer's protocol. Two days after transfection, medium was removed and replaced with fresh CDI maintenance media. Recordings of cardiac field potential recordings, contractility, and $Ca^{2+}$ transients were collected from cardiomyocyte monolayers at 7 days post siRNA silencing.

For transduction of human iPSC-CM isogenic lines with TN-401, cells were thawed and recovered in iCell Cardiomyocyte Plating medium (FUJIFILM Cellular Dynamics), and brought to a cell density of 200,000 cells/mL or 625,000 cells/mL for a 96 well or 384 well plate format, respectively. An appropriate volume of TN-401 for multiple MOIs was added to the cells seeded on pre-coated Matrigel plates at a density of 300,000, 40,000 or 15,000 cells per well for 24, 96 or 384 well plate format, respectively. Following overnight incubation, medium was replaced by fresh metabolic maturation medium. For cells seeded on MEA plates, a density of 16,000 cells/well were used.

### Small Molecule Treatment

Between days 6 and 12 post-seeding of monolayers or 21 days after casting for EHTs, iPSC-CM monolayers and EHTs were treated with small molecule inhibitors or enhancers at the indicated final concentrations in the table below. Small molecules were prepared as working stock solutions in DMSO and diluted to their respective final concentrations in the culture medium. Vehicle control groups were treated with DMSO at ≤ 0.1% of the volume of the culture medium. Small molecules were refreshed at 24 hours post initial treatment. Measurements for electrophysiology, calcium transients, and contraction were collected at 1, 24, and 48 hours.

| Vehicle or Compounds | [Working Stock] | [Final] | µl/10 ml |
|---|---|---|---|
| DMSO | / | / | ≤ 10 |
| Etomoxir | 10 mM | 4 µM | 4 |

| | | | |
|---|---|---|---|
| Dgat1i (AZD7687) | 10 mM | 5 µM | 5 |
| Bezafibrate | 100 mM | 100 µM | 10 |
| Seladelpar | 100 mM | 10 µM | 1 |
| Isoproterenol (DMSO) | 10 mM | 10 µM | 10 |
| Phenformin (H2O) | 100 mM | 80 µM | 8 |

### Glucose and lactate measurements

After two weeks of culture in 96-well Seahorse plates, the glucose, lactate, sodium and potassium levels in human iPSC-CM cell culture supernatant was determined by a BioProfile FLEX2 with Chemistry MicroSensor card.  The day before FLEX2 measurements, media was removed from all wells and replaced with twice the working volume of fresh culture media (200µL Mercola Media per well) and cells were incubated for 15 minutes to equilibrate the media.  After 15 minutes, one working volume of cell culture supernatant was removed (100µL per well) from each well and stored frozen at -80°C for use as the initial time point.  The cells were incubated overnight in the remaining cell culture media (100µL per well).  The next day, the remaining cell culture media was harvested from each well for use as a final time point. All samples were stored frozen at -80°C until the time of measurement.  Samples were thawed at room temperature immediately before measurement and vortexed briefly to ensure homogeneity.  Media from three wells were pooled (3 x 100µL) to generate a single analytical sample of sufficient volume to be injected on the FLEX2 system.  The same wells were pooled for initial samples and final samples.  To correct for evaporation during incubation between initial and final time points, the measured glucose and lactate concentrations from each analytical sample were normalized by an average of the measured sodium and potassium concentrations from the same sample.  After normalizing, final timepoints were divided by initial timepoints to calculate a "percent of initial" value for glucose and lactate for each pooled analytical sample.  Two or three separate pooled analytical samples per treatment group were used for statistical analysis.

### Seahorse analysis

The Seahorse XFe96 extracellular flux analyzer was used to measure human iPSC-CM bioenergetics. Cardiomyocytes were thawed and seeded onto plates, coated with Matrigel, at a target density of 18,000 (CDI CM[2]) or 20,000 (in-house differentiated isogenic lines) cells per well. The day of analysis, Seahorse FluxPaks were calibrated on the XFe96. Then, cell culture medium was exchanged for Agilent Seahorse XF DMEM Basal Medium supplemented with 2 mM glutamine, 10 mM glucose, BSA-palmitate (100µM), and 0.1 mM sodium pyruvate and immediately transferred to the XFe96. Cells were equilibrated in the Seahorse instrument for 20 minutes before initial measurements. For the Seahorse assays, inhibitors were injected during the measurements as follows; MitoStress Test, oligomycin (2 µM), FCCP (1 µM), rotenone and antimycin A (1 µM, each); Palmitate Oxidation Stress Test, etomoxir (4 µM) or vehicle, oligomycin (2 µM), FCCP (1 µM), rotenone and antimycin A (1 µM, each); ATP Rate Assay, oligomycin (2 µM), rotenone and antimycin A (1 µM, each).

The oxygen consumption rate (OCR) and extracellular acidification rate (ECAR) values were normalized to the number of nuclei per well, quantified by Hoechst 33342 staining. Basal OCR was defined as the difference between the time point before oligomycin (MitoStress Test) or etomoxir (Palmitate Oxidation Stress Test) and the minimum OCR after rotenone/antimycin A. Maximal OCR was the OCR difference between maximum OCR after uncoupler FCCP and minimum OCR after rotenone/antimycin A.

ATP Rates were calculated from OCR and ECAR as described in the Agilent ATP Rate Assay kit brochures and white papers. Briefly, ATP-linked OCR was defined as the difference between OCR and the average post-oligomycin OCR. ATP-linked OCR (pmol $O_2$/min) was converted to mitoATP Rate (pmol ATP/min) by multiplying by 5.5; assuming 2 oxygen atoms/$O_2$ molecule and 2.75 ADP molecules phosphorylated per oxygen atom consumed. Separately, ATP-linked OCR was converted to a mitochondrial proton efflux rate (mitoPER; pmol $H^+$/min) by multiplying by 0.61; assuming the contribution of mitochondrial derived $CO_2$ to extracellular acidification equals 0.61 $H^+/O_2$ consumed (empirically determined by Agilent for multiple cell lines in XFe96 plates). ECAR (mpH/min) was converted to total proton efflux rate ($PER_{Total}$; pmol $H^+$/min) by multiplying by 8.76; combining the Seahorse Assay Buffer factor empirically determined to be 2.4 mmol $H^+$/L/pH for the buffer used in these studies, the XFe96 microchamber volume of 2.28 µL/well, and the $K_{vol}$ scaling factor of 1.6 (empirically determined by Agilent to be the XFe96 effective microchamber volume / the expected microchamber volume). The proton efflux rate from glycolysis (glycoPER; pmol $H^+$/min) was defined as the difference between $PER_{total}$ and mitoPER. Lastly, glycoPER is converted to glycoATP Rate by multiplying by 1; assuming 1 ATP molecule was produced from glycolysis for every 1 effluxed $H^+$ derived from glycolysis. After calculations, mitoATP Rates and glycoATP Rates were reported in units of pmol ATP/min and normalized by nuclear count.

### Fatty acid uptake and storage assays

Four days prior to the assay, 20,000 human iPSC-CMs were seeded on a Matrigel coated 96-well plate. The fatty acid was evaluated with the QBT Fatty Acid Uptake Assay (Molecular Devices) that uses a BODIPY®-dodecanoic acid fluorescent fatty acid analog coupled with a proprietary quench technology. Cells were washed 3x with serum-free media and preincubated in serum-free media for 2 hours. A baseline measurement was preformed prior to the addition of the fluorescent fatty acid mixture, and every 10 mins after the addition by using a Cytation 5 Cell Imaging Multimode Reader (Agilent BioTek) at 485/528 nm.

LipidSpot 610 lipid droplet stain (Biotium) are fluorogenic neutral lipid stains that rapidly accumulate in lipid droplets, where they become brightly fluorescent.

### Evaluation of contractility, calcium transients, and electrophysiological properties of iPSC-CMs

*Contractility analysis of monolayers*

Human iPSC-CMs were seeded on pre-coated Matrigel plates at a density of 20,000 cells per well for 384-well plate or 20,000 cells of CDI CM$^2$ (FUJIFILM Cellular Dynamics) or 40,000 cells of in-house differentiated isogenic iPSC-CMs per well for 96-well plate. Contractility of iPSC-CMs was measured in bright field by SONY SI8000 Cell Motion Imaging System (Sony Biotechnology). Eight seconds videos of 300 frames at

38 fps were recorded. Data were processed and analyzed using the CuriBio Pulse data analysis platform[10] (CuriBio).

*Electrophysiological Evaluation of Monolayers*

The microelectrode array in the center of each well of the CytoView MEA 24-well plates (Axion Biosystems) were coated/spotted with 10 µL droplet of Matrigel diluted at a 1:100 ratio in KnockOut DMEM (Gibco) and incubated for 4 hours at 37°C with 5% $CO_2$ in a cell culture incubator. To increase humidity of the plate environment and minimize evaporation from the wells, 10 mL of sterile UltraPure Distilled Water (Invitrogen) was added to the on-plate reservoirs located around the perimeter and between wells of the plates. Subsequently, Matrigel was removed, and human iPSC-CM droplets were plated on top of the microelectrode array at a density of 16,000 cells/well in 8 µL of iCell cardiomyocytes plating medium. The cells were maintained in a cell culture incubator for 2-4 hours to allow cells to settle and adhere to the microelectrode array, and thereafter, 0.5 mL of iCell Cardiomyocytes maintenance medium was added to each well. A full volume media change (maturation media) occurred 24-hour post seeding and media change continued every other day until the end of the experimental period.

Cardiac field potential recordings in iPSC-CM monolayers were performed using the Maestro Pro MEA platform (Axion Biosystems) at 37°C and 5% $CO_2$. Real-time data on field potential duration (FPD), beat rate, spike amplitude, spike slope and conduction velocity were collected simultaneously across all electrodes (16 per well). Analysis was performed using AxIS Navigator software and the Cardiac Analysis Tool[11] (Axion BioSystems).

*$Ca^{2+}$ transient recordings of Monolayers*

The bottom of Falcon 24-well clear flat bottom cell culture plates (Corning) were coated with 1 mL of Matrigel diluted at a 1:100 ratio in KnockOut DMEM and incubated overnight at 37°C with 5% $CO_2$ in a cell culture incubator. Next day, Matrigel was removed, and human iPSC-CM were thawed and plated at a density of 300,000 cells/well in 1 mL of iCell cardiomyocytes plating medium. The cells were maintained in a cell culture incubator for 4 hours to allow cells to settle and adhere to the Plate, and thereafter, media in each well was replaced with 1 mL of iCell Cardiomyocytes maintenance medium. A full volume maturation media change occurred every other day post seeding until the end of the experimental period.

Calcium transient recordings in human iPSC-CM monolayers were performed using the CuriBio Nautilai. Monolayers were first incubated (37°C with 5% $CO_2$) with 5 µM Fluo-4 AM (Invitrogen) in 1.2 mM $Ca^{2+}$ Tyrode's solution (Boston BioProducts) for 30 min. Fluo-4 AM was prepared by dissolving it in DMSO (Sigma Aldrich) with 10% of Pluronic F-127 (Invitrogen). Thereafter, the dye was removed and cells incubated (37°C with 5% $CO_2$) in 1.2 mM $Ca^{2+}$ Tyrode's solution for 10 min. Cells were then stabilized by electrical stimulation at 1 Hz or 1.5Hz (Voltage: 10 Volts and Pulse width: 3 ms) for 5 minutes. Optical real time $Ca^{2+}$ transient data acquisition with the Nautilai was performed (LED intensity: 20%, Frames Per Second: 250, and Duration 20- 30 seconds). Analysis was performed using Nautilai cloud data analysis platform Pulse (CuriBio).

### Evaluation of contractility properties and calcium handling of EHT

Contractility measurements were conducted using the Mantarray Instrument (CuriBio) at 37$^o$C and 5% $CO_2$. One-minute videos were recorded and subsequently processed with the CuriBio Pulse data analysis platform (CuriBio).

For calcium handling measurements, a 1.2 mM $Ca^{2+}$ (Sigma) Tyrode's Solution (Modified II, pH = 7.4) (Boston BioProducts) was prepared. Fluo-4 AM green fluorescent calcium dye (Invitrogen) was diluted to a 1 mg/mL solution in DMSO with 10% Pluronic F-127, and mixed with the 1.2 mM $Ca^{2+}$ Tyrode's Solution at a final concentration of 4.5 µM. EHTs were loaded with Fluo-4 AM in Tyrode's Solution and incubated for 30 min at 37$^o$C, 5% $CO_2$. Fluoa-4 AM was removed and EHTs were incubated for 10 min at 37$^o$C, 5% $CO_2$ in 1.2 mM $Ca^{2+}$ Tyrode's Solution (no Fluo-4 AM). EHTs were then imaged with Nautilai. Calcium transients were recorded for 30 s at 250 fps at 37$^o$C, and data were processed and analyzed using the CuriBio Pulse data analysis platform.

### Immunofluorescence imaging and quantification of iPSC-CMs

Human iPSC-CMs were fixed in 4% paraformaldehyde (Thermo Fisher Scientific) for 15 min and permeabilized with PBS + 0.1% Triton-X100 (Millipore) (PBST) at room temperature for 15 min. Cells were washed with PBS three times followed by blocking with PBST + 4% bovine serum albumin (BSA) (Thermo Fisher Scientific) for 1 h. Cells then were incubated with primary antibodies against PKP2 (Invitrogen, rabbit polyclonal PA5-53144 or rabbit polyclonal PA5-66048) at 1:200 dilutions, DSP (Invitrogen, rabbit polyclonal 25318-1-AP; Sigma, mouse monoclonal MABT1492) at 1:100 dilution or PKG (Sigma, mouse monoclonal P8087 or rabbit polyclonal SAB4501616) at 1:200 dilutions in PBS + 4% BSA for overnight. After washing with PBS three times, cells were then incubated with donkey anti-mouse or anti-rabbit Alexa Fluor 488, 594 or 647 (Invitrogen) at 1:500 dilutions in PBS + 4% BSA for 2 h. Cells were washed 4 times and counterstained with 10 mg/mL Hoechst 33342 (Thermo Fisher Scientific) for 15 min, imaged, and acquired on either Cytation 5 Cell Imaging Multimode Reader or Molecular Devices ImageXpress Micro Confocal High-Content Imaging System (Molecular Devices). Confocal images were acquired from 9 independent cell areas for each well in a format of 96-well plates. Images in different fluorescence channels were merged using ImageJ software and analyzed using CellProfiler, or directly analyzed with Cytation 5 analysis software (Agilent BioTek).

### Statistics and reproducibility

The numbers of technical and biological replicates and animals for each experiment are indicated in the figure legends. Normality and lognormality tests were performed first to determine whether a dataset is normally distributed. Statistical analyses, ordinary One-Way ANOVA (Tukey’s post-hoc test), ordinary Two-Way ANOVA (Sidak's post-hoc test), and Student’s t-test, were performed using GraphPad Prism 9. Significant differences were defined as $p < 0.05$. Error bars in all mouse studies represent SEM (Standard Error of the Mean). Error bars in all cell biology studies represent SD (Standard Deviation). Statistical tests for each individual experiment are provided in the figure legend.

Statistical analyses for RNA sequencing data are detailed below.

### Transcriptional analysis by RNA sequencing

From each replicate, 100 ng total RNA was extracted via the polyA-tail-specific protocol according to Illumina Inc. RNA quality control was performed before library preparation using Agilent TapeStation instrument. The RNA libraries were prepared using a Stranded Total RNA Library Prep with Ribo-Zero Plus kit (Illumina), which also removes ribosomal RNA. The libraries were sequenced as 2x50 base pair paired-end reads using Illumina NovaSeq 6000 using V1.5 reagent kit on S1 flow cell with an average of 25.66 million reads per each read file (51.32M reads per sample). After adapter trimming by fastp (version 0.23.3), raw RNA-seq reads from mouse hearts in fastq format were aligned with Salmon (version 1.8.0) to the GENCODE (version M30, July 2022) reference transcript assembly (GRCm39 and Ensembl 107) using best practice parameters to ensure mapping validity and reproducibility (--seqBias --gcBias --posBias --useVBOpt --rangeFactorizationBins 4 --validateMappings --mimicStrictBT2). Next, a script using R package *tximport* was used to generate an expression matrix normalized to transcripts per million (TPM). In this analysis, we only used genes detected in at least 10% of all samples. Protein-coding genes were determined using Ensembl release *Mus musculus* annotations (GRCm39, July 2022) and extracted by *biomaRt* (version 2.52.0). Mitochondrial genes were also omitted, followed by renormalization to TPM. These gene expression values were then log2-transformed after addition of 1 as pseudo-count. Expression patterns of key genes associated with functions of interest were visualized across treatment groups with boxplots generated using the *ggplot2* R package. Expression values from both left and right ventricles were included in the boxplots. Relative gene expression levels across groups and two ventricles are also presented in scaled values per gene in the heatmaps. Heatmaps were generated in R using *ComplexHeatmap* package.

For initial assessment and identifying presence of cluster patterns in the transcriptome, Principal Component Analysis (PCA) models were generated in R using the `prcomp` function from the *stats* package. The first two principal components were used to visualize group level differences across samples in a PCA plot generated using *ggplot2* and *ggfortify* packages with the `autoplot` function. Differential gene expression analysis was then performed by comparing each two groups of interest using Welch's *t*-test on pseudo-log normalized TPM values. The obtained *t* statistics values were used to rank-order the genes for the downstream functional analyses. Volcano plots were then generated to visualize the top positive and negative differentially expressed genes (DEGs) using the *ggplot2* R package. The top DEGs are the set of genes with the highest and lowest t-statistics values. To evaluate functional effects, we performed Gene Set Enrichment Analysis (GSEA)[12] on the gene list pre-ranked by t-statistics obtained from differential gene expression analysis, using the *clusterProfiler* R package. GSEA assesses whether differences in expression of predefined gene sets between two phenotypes are concordant and statistically significant. Gene sets were obtained from positional, curated canonical pathways, transcription factor targets, Gene Ontology, cell type signatures and Hallmark collections in Human MSigDB (v2023.1.Hs)[12,13,14]. Upon performing GSEA, these gene sets were only considered statistically significant if the false discovery rate (Q value) was less than 0.25 as determined with multiple hypothesis testing correction using the BH-correction method[15]. The normalized enrichment score, which reflects the degree to which a gene set is overrepresented in the ranked list and normalized for gene set size, was used to select significantly altered gene sets. Trends in normalized enrichment scores for some gene sets of interest were shown in heatmaps, which were generated in R using *ComplexHeatmap*.
